# Discovery of active mouse, plant and fungal cytochrome P450s in endogenous proteomes and upon expression *in planta*

**DOI:** 10.1101/2024.03.18.585456

**Authors:** Maria Font Farre, Daniel Brown, Reka Toth, Chidambareswaren Mahadevan, Melissa Brazier-Hicks, Kyoko Morimoto, Farnusch Kaschani, John Sinclair, Richard Dale, Samantha Hall, Melloney Morris, Markus Kaiser, Aaron T. Wright, Jonathan Burton, Renier A. L. van der Hoorn

## Abstract

Eukaryotes produce a large number of cytochrome P450s that mediate the synthesis and degradation of diverse endogenous and exogenous metabolites. Yet, most of these P450s are uncharacterized and global tools to study these challenging, membrane-resident enzymes remain to be exploited. Here, we applied activity profiling of plant, mouse and fungal P450s with chemical probes that become reactive when oxidized by P450 enzymes. Identification by mass spectrometry revealed labeling of a wide range of active P450s, including six plant P450s, 40 mouse P450s and 13 P450s of the fungal wheat pathogen *Zymoseptoria tritici*. We next used transient expression of GFP-tagged P450s by agroinfiltration to show ER-targeting and NADPH-dependent, activity-based labeling of plant, mouse and fungal P450s. Both global profiling and transient expression can be used to detect a broad range of active P450s to study e.g. their regulation and discover selective inhibitors.

## Introduction

Cytochrome P450s (P450s) are numerous in eukaryotes and play crucial, diverse roles in the metabolism of endogenous and exogenous metabolites. P450s are thiolate-ligated haem monooxygenases able to oxidize a broad range of small molecules. The vast majority of eukaryotic P450s are membrane bound proteins, mostly integrated in the membrane of the endoplasmic reticulum (ER), with the catalytic domain facing the cytosol [1,2]. Nearly all P450s require nicotinamine adenine dinucleotides (NADH/NADPH) as cofactors delivering reductants. Cytochrome P450 reductases (CPRs) are the common redox partner in microsomal P450 systems and are responsible for the transfer of reducing equivalents from NADPH to P450s [3,4].

Despite the importance of P450s in metabolism of endogenous and exogenous compounds, these enzymes are notoriously challenging to study for several reasons. First, most eukaryotes encode large numbers of P450s, e.g. 102 in mice [5] and 244 in model plant *Arabidopsis thaliana* [6]. Second, their activity depends on reductants produced by CPRs. Third, they are integral membrane proteins which usually require the isolation and testing of microsomal fractions. Finally, for most P450s, no specific substrates or inhibitors are known or available. More general and comprehensive approaches to study P450s are needed to characterize these important enzymes. Activity-based protein profiling (ABPP) is a powerful method to monitor, identify and characterize active enzymes of different families using chemical probes that label these enzymes in an activity-dependent manner [7–9]. These chemical probes have a reactive group (warhead) that makes a covalent bond with the target enzyme, and a reporter tag (e.g. biotin or fluorophore) or an alkyne minitag to be coupled to a reporter tag via ‘click chemistry’, which is used for the purification and detection of labeled proteins [10].

Previously, activity-based probes were introduced to study specific P450s in mammals [11–12], pyrethroid-metabolizing P450s in insects [13] and ammonia monooxygenase in bacteria [14]. These P450 probes react as suicide substrates because they are substrate-like molecules that become reactive upon mono-oxygenation by P450s and subsequently react with nucleophiles within the substrate pocket of P450s. Therefore, although labeling is strictly dependent on the activity of the P450, labeling does not necessarily occur at active site residues, in contrast to activity-based probes targeting other enzyme families [8]. In this work, we used P450 probes to discover active P450s during transient expression in agroinfiltrated leaves, which are used for metabolic pathway reconstruction, furthermore in *Zymoseptoria tritici*, an important fungal wheat pathogen that uses P450s to detoxify fungicides. We also used plant-based transient expression to display specific active P450s from mouse and fungi and used this to demonstrate that the P450s have different sensitivities for the probes, indicating that probe cocktails can be used to broaden P450 profiling.

## Results

Wright and coworkers designed various activity-based probes to label human P450s [11]. These probes have not been tested on P450s of non-metazoans. We resynthesized five different P450 probes (**Figure 1a**) to investigate P450 labeling in different organisms. The reactive groups in these probes are aryl alkynes (DB086, DB088, DB089 and DB096) or furan (DB080). These probes react with P450s because they mimic a P450 substrate and are oxidized by P450s, resulting in a reactive intermediate that quickly reacts with nearby residues within the substrate binding pocket of P450s, resulting in a covalent, irreversible, activity-based labeling of these enzymes (**Figure 1b**, [11]). The P450 probes also carry an alkyne minitag that can be coupled to azide-biotin or azide-fluorophore via click chemistry for purification or fluorescence scanning, respectively. In this work, we frequently use picolyl-azides for more efficient click chemistry reactions (**Figure 1c**).

**Figure 1.**
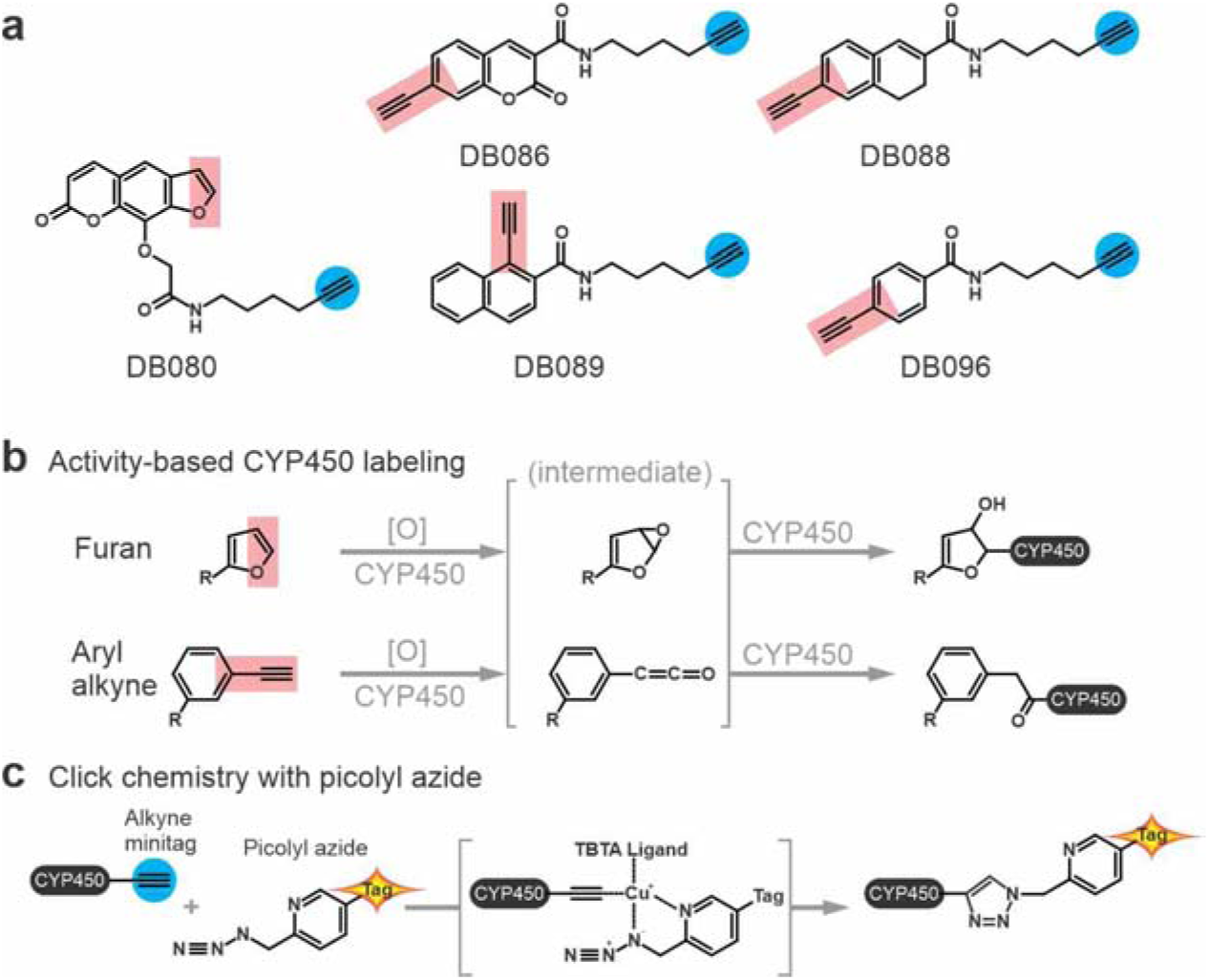
P450 probes and reaction mechanisms. **(a)** Structures of five activity-based probes for P450s. All probes contain an alkyne minitag (right, blue circles), for coupling to azide-fluorophores or azide-biotin via click chemistry. The reactive group (left, red rectangles) is an aryl alkyne-based reactivity group, except for DB080, which contains a furan reactivity group. **(b)** Labeling mechanisms. The P450 probes are oxygenated by P450, leading to a reactive intermediate that covalently links to nearby residues in the substrate binding site of P450s. **(c)** Click chemistry with picolyl azide. The coupling of the P450-alkyne to azide-tag is catalyzed by copper(I), which is stabilized by the ligand (tris((1-benzyl-1H-1,2,3-triazol-4-yl) methyl)amine, TBTA), the picolyl, and the azide, resulting in a stable triazole linker between the P450 protein and a reporter tag. The reporter tag is either biotin or a fluorophore.

### ZmCYP81A9 is inhibited by P450 probes

To test if the probes can also inhibit plant P450s, we monitored inhibition of maize (*Zea mays*) *Zm*CYP81A9, for which activity assays have been established, because it is an important herbicide-metabolizing enzyme responsible for the modification of herbicides with unrelated modes of action [15–17]. *Zm*CYP81A9 was expressed in brewer’s yeast (*Saccharomyces cerevisiae*, [18]) and microsomes from this strain were isolated and used to monitor the conversion of bentazone, leading to hydroxybentazone (**Figure 2a**), which can be measured by LC-MS [19]. *Zm*CYP81A9 activity can be inhibited with the broad-range P450 inhibitor 1-aminobenzotriazole (1-ABT, [20]), which was included as a positive control for inhibition. HPLC analysis revealed that, besides 1-ABT, all tested P450 probes could inhibit bentazone hydroxylation by *Zm*CYP81A9 (**Figure 2b**).

**Figure 2.**
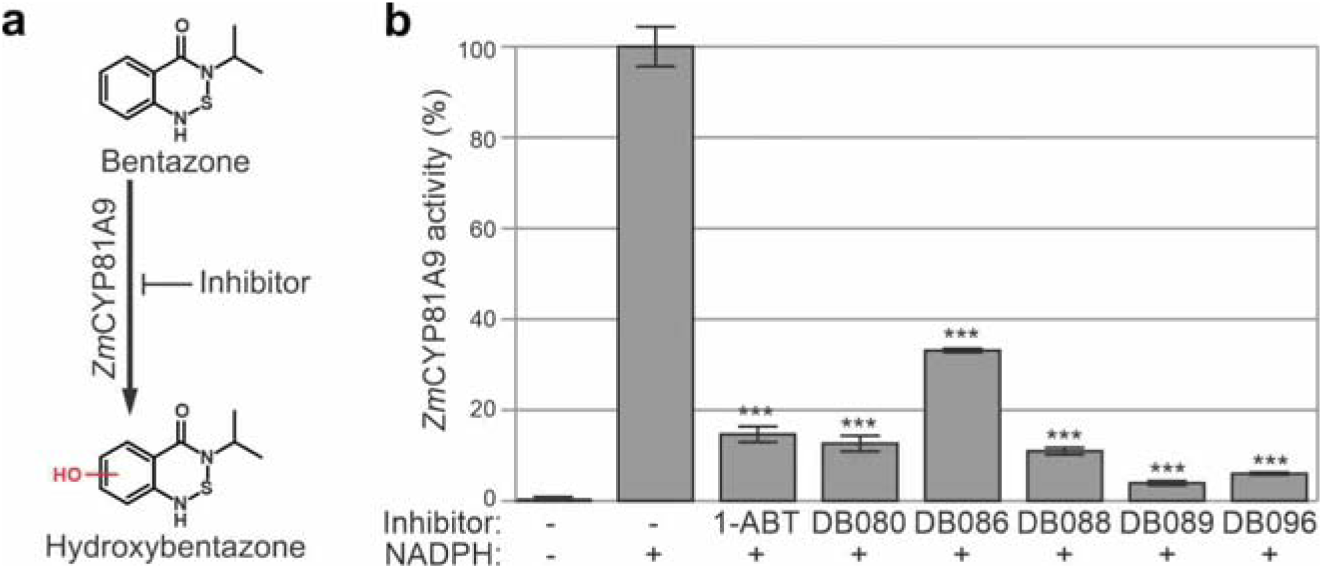
Maize *Zm*CYP81A9 is inhibited by P450 probes. **(a)** Reaction scheme of hydroxylation of bentazone, catalyzed by *Zm*CYP81A9. **(b)** P450 probes inhibit *Zm*CYP81A9 activity. Microsomes isolated from yeast expressing *Zm*CYP81A9 were preincubated with 300 μM or without inhibitor and with or without 1 mM NADPH for 20 minutes at 28 °C, and then incubated for 120 minutes with 200 μM bentazone at 28 °C. Metabolites were extracted with acetonitrile and analyzed by LC-MS to measure peak areas of hydroxylated bentazone. The no-inhibitor-control was set at 100% activity and used to calculate the relative activity and statistical relevance of the other samples (t-test; ***, P<0.001). Error bars represent standard deviation of n=3 replicates.

### Transiently expressed ZmCYP81A9 accumulates in the ER and can be labeled with P450 probes

Having established that *Zm*CYP81A9 can be inhibited by the probes, we produced *Zm*CYP81A9 to study its labeling upon transient expression in *Nicotiana benthamiana* leaves by agroinfiltration. Agroinfiltration is a quick and easy way to overexpress proteins in a eukaryotic host, and is based on the infiltration of *Agrobacterium tumefaciens* delivering transfer DNA with genes of interest into plant cells (**Figure 3a**). *Zm*CYP81A9 was C-terminally tagged with green fluorescent protein (GFP, **Figure 3a**) to enable quick verification of protein accumulation by GFP fluorescence [21] and to shift the molecular weight of the P450 away from the 50 kDa region that contains endogenous P450s and is co-occupied by the large subunit of Ribulose-1,5-bisphosphate carboxylase/oxygenase (RBCL), an abundant plant protein that causes background labeling (see below).

**Figure 3.**
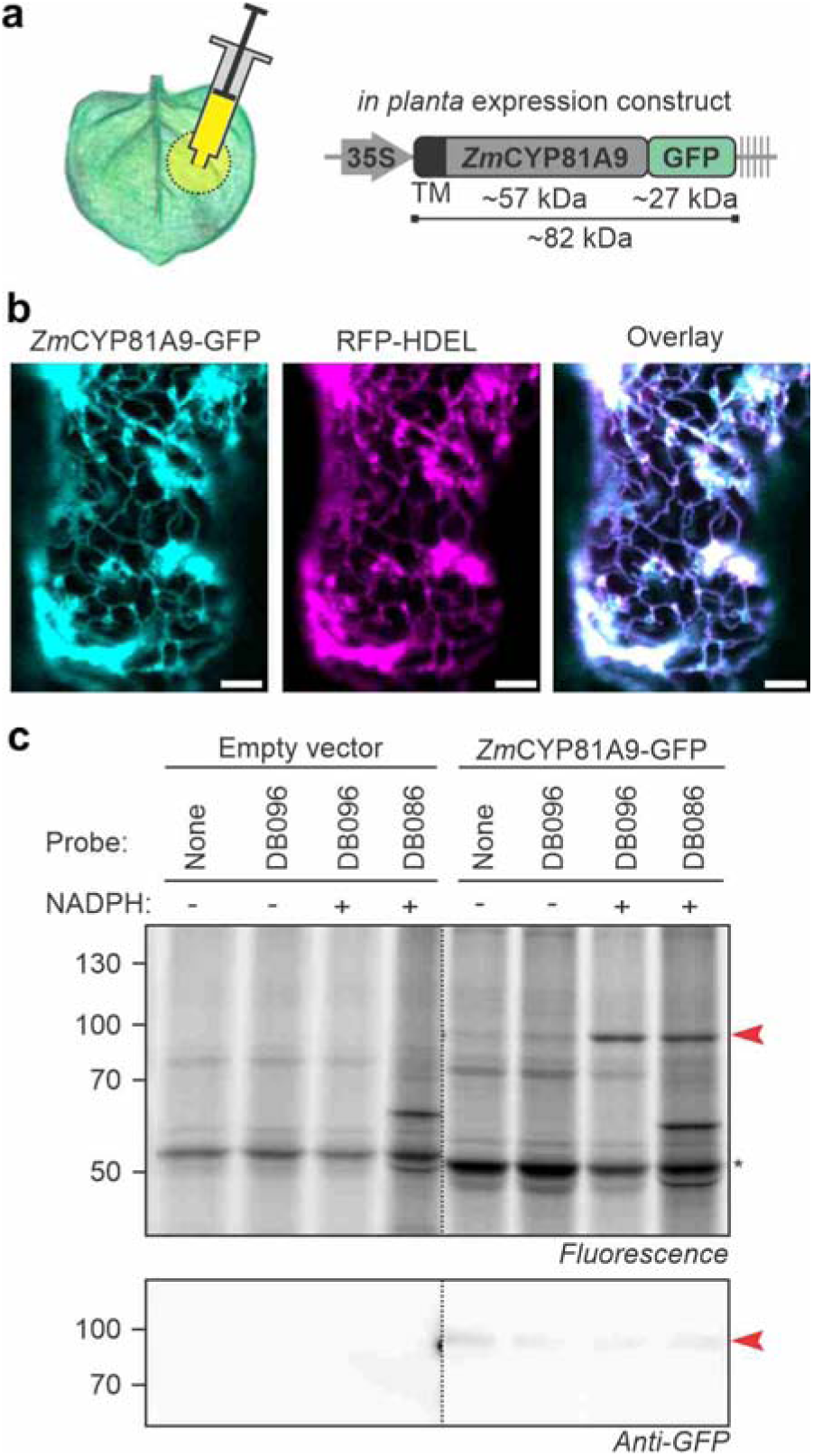
Transiently expressed Maize *Zm*CYP81A9 is labeled by P450 probes. **(a)** Transient expression of GFP-tagged ZmCYP81A9 by agroinfiltration of *Nicotiana benthamiana*. **(b)** *Zm*CYP81A9-GFP localizes to the ER. *Zm*CYP81A9-GFP was transiently co-expressed with ER marker RFP-HDEL by agroinfiltration. Confocal images were taken 5 days after agroinfiltration. Scalebar 5 μm. Experiment was repeated twice with similar results. **(c)** *Zm*CYP81A9-GFP is labeled by the P450 probes. Microsomes were isolated from agroinfiltrated leaves expressing *Zm*CYP81A9-GFP and labeled for 1.5hr with various probes (10 μM) with or without 1 mM NADPH. Labeled proteins were coupled to Cy5-picolyl-azide via click chemistry and visualized by in-gel fluorescence scanning. Western blot with anti-GFP antibodies shows accumulation of *Zm*CYP81A9-GFP at its expected MW. *, RBCL.

To investigate if *Zm*CYP81A9-GFP accumulates in the ER membrane in plants, we performed confocal microscopy on leaves transiently co-expressing *Zm*CYP81A9-GFP with ER marker protein RFP-HDEL [22] to confirm its localisation in the ER. Confocal imaging demonstrates that *Zm*CYP81A9 localizes in a subcellular structure that is typical for the ER network (**Figure 3b**). This network showed a high correlation between *Zm*CYP81A9-GFP and RFP-HDEL fluorescence (Pearson’s correlation = 0.886±0.099, Supplemental **Figure S1a**). Thus, *Zm*CYP81A9-GFP correctly localizes in the ER membrane upon transient expression *in planta*.

We next used microsomes isolated from agroinfiltrated leaves expressing *Zm*CYP81A9-GFP for labeling with the different P450 probes. This experiment revealed an NADPH-dependent fluorescent signal for DB086 and DB096 (**Figure 3c**), indicating that *Zm*CYP81A9-GFP is labeled with these P450 probes. This labeling experiment also displays a strong 50 kDa signal that is NADPH-independent and also appears in the click control sample (**Figure 3c**). This protein is very likely RBCL, the most abundant protein in leaf extracts. This strong background hinders the detection of labeling of endogenous P450s, which all have predicted molecular weights of 50-55 kDa. Thus, tagging of heterologous P450 with GFP is not only useful to quickly detect their accumulation; it also shifts the labeled protein away from the size of endogenous P450s and the background labeling of RBCL.

### Maize and N. benthamiana P450s are labeled with P450 probes

We next aimed to confirm *Zm*CYP81A9-GFP labeling by proteomics on purified labeled proteins. Microsomes from agroinfiltrated leaves expressing *Zm*CYP81A9-GFP were labeled with DB088 and DB096 and labeled proteins were biotinylated and purified on streptavidin beads. The beads were incubated with trypsin/Lys-C and the released peptides were analysed by LC-MS/MS for n=3 biological replicates. Click control (CC) samples were included as a negative control that were treated the same way except that no DB088/DB096 was added. Volcano plots showed *Zm*CYP81A9-GFP as the top hit in both pull-down experiments (**Figure 4a**, Supplemental **File S1-S2**), confirming that *Zm*CYP81A9-GFP is labeled by both DB088 and DB096. Interestingly, we also detected five endogenous P450s of *Nicotiana benthamiana* amongst the enriched proteins with DB088 labeling, and four of these also upon DB096 labeling (**Figure 4a**), indicating that these endogenous P450s are also labeled by P450 probes.

**Figure 4.**
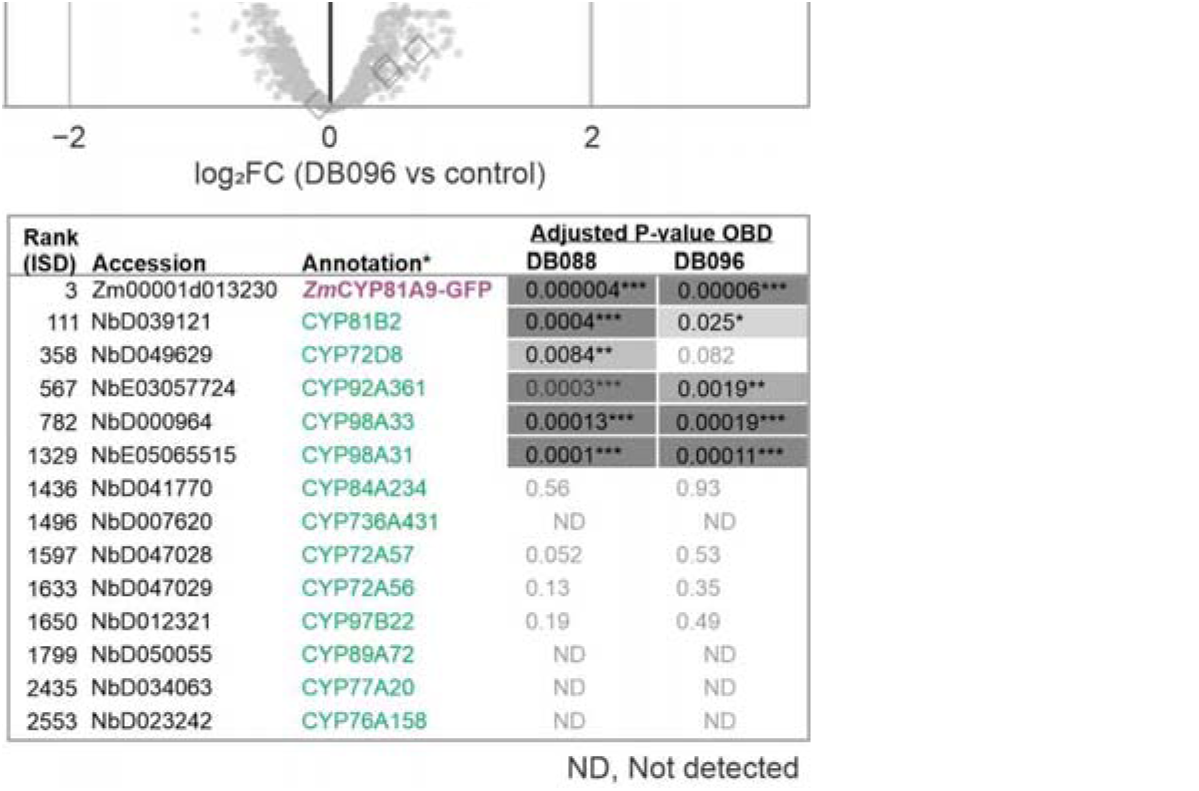
Maize *Zm*CYP81A9 and *N. benthamiana* P450s are labeled with P450 probes. **(a)** Volcano plots for detection of labeled proteins purified from microsomes of *Zm*CYP81A9-GFP-expressing leaves incubated with DB088 (left) or DB096 (right). Microsomes isolated from agroinfiltrated leaves expressing *Zm*CYP81A9-GFP were labeled with and without 10 μM probes in the presence of 1 mM NADPH for 1.5 hr. Labeled proteins were coupled to azide-biotin using click chemistry and biotinylated proteins were purified on streptavidin beads, digested with trypsin (on-bead-digest, OBD) and released peptides were identified by MS. The log2 fold change (log2FC) was plotted against the significance (-log10FDR) for n=3 biological replicates. Highlighted are proteins that are significantly more (green) or less (red) abundant in the probe labeled samples when compared to the click-chemistry control, showing P450s as diamonds. **(b)** Shotgun proteomics of the input sample. Microsomes isolated from agroinfiltrated leaves expressing *Zm*CYP81A9-GFP were digested with trypsin (in-solution-digest, ISD) and analyzed by MS. Proteins were ranked on their total MS spectral intensity and the P450s are highlighted. Identified P450s and their rank in the ISD dataset and the log2 of the MS intensity, followed by the adjusted P-values in on-bead digest analysis presented in (a). *, the annotation of *N. benthamiana* P450s is based on the closest homology.

Finally, to investigate the relative abundance of the detected P450s, we analysed the proteomes of the input microsomes by in-solution-digest with trypsin/Lys-C. We ranked these by their total spectral intensities, which is a good estimate of relative protein abundance [23–24]. When ranked on total MS intensity, *Zm*CYP81A9-GFP was found on rank-3 (**Figure 4b**, Supplemental **File S3**), consistent with being overexpressed by agroinfiltration, and being the most robustly detected P450 in labeling experiments. We detect another 13 P450s of *N. benthamiana* in these samples (**Figure 4b**). Of these endogenous P450s, *Nb*CYP81B2 has the highest protein intensity, at rank-111. Notably, only the five most abundant *N. benthamiana* P450s in the microsomes of agroinfiltrated leaves were significantly enriched upon labeling (**Figure 4b**), indicating that detection of P450s by both DB088 and DB096 correlates with protein abundance.

### Effective P450 labeling in mouse liver microsomes

The absence of NADPH-dependent labeling of endogenous P450s prompted us to perform labeling experiments on mouse liver extracts, which were previously used to established P450 profiling [12]. All five P450 probes caused NADPH-dependent labeling, resulting in several signals of 50-60 kDa, consistent with the predicted molecular weight (MW) of mouse P450s (**Figure 5a**). Strongest labeling was observed for DB088, DB096 and DB143 probes.

**Figure 5.**
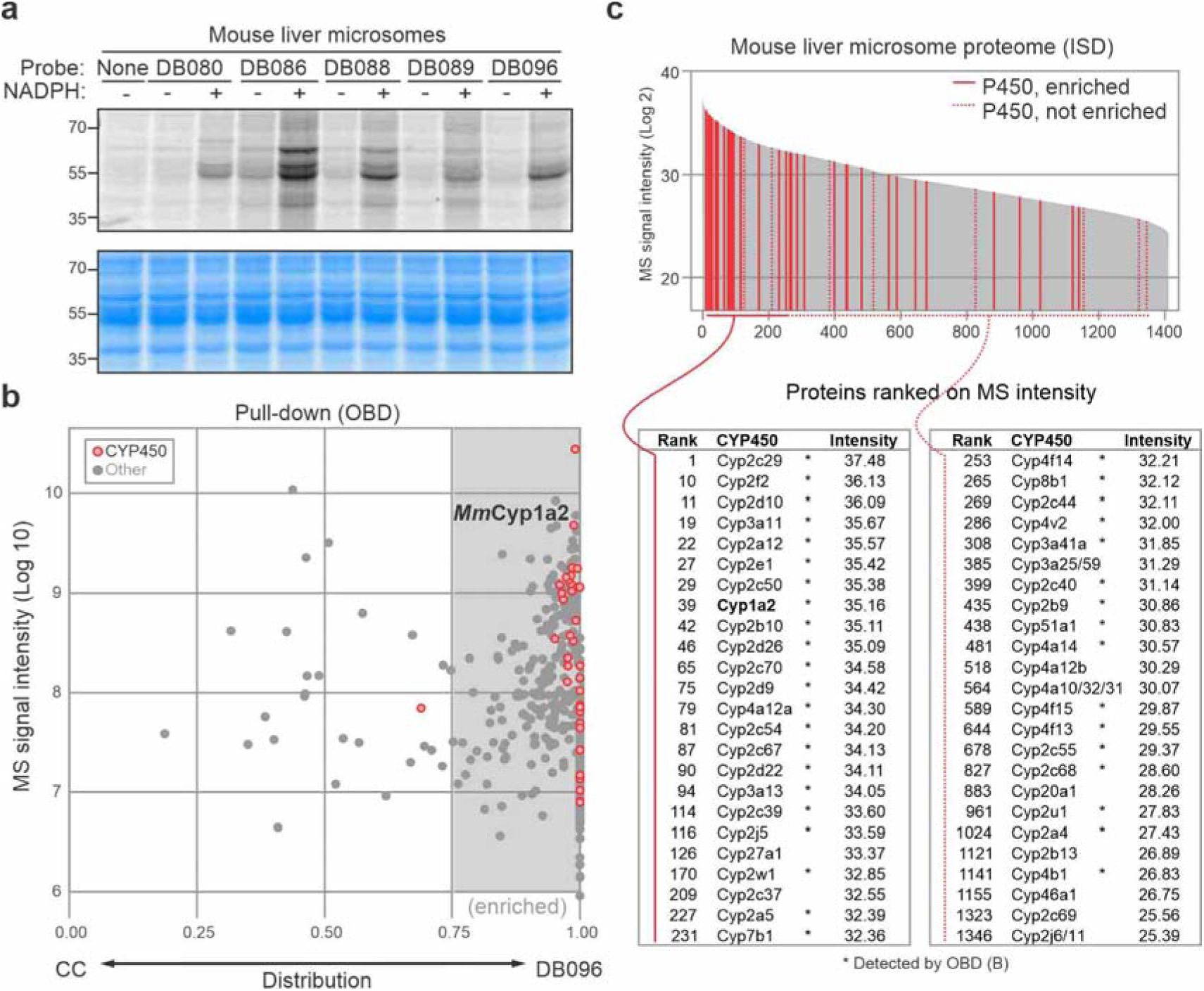
Labeling of P450s in mouse liver microsomes. **(a)** NADPH-dependent labeling in mouse liver microsomes. Microsomes were labeled for 1.5 hr with 20 μM probe with and without 1 mM NADPH. Labeled proteins were coupled to Cy5-picolyl-azide via click chemistry, separated on SDS-PAGE gels and visualized by in-gel fluorescence scanning. **(b)** Distribution graph summarizing proteins identified upon purification of labeled proteins from mouse liver microsomes. Microsomes were incubated for 1.5hr with 30 μM DB096 or an equal volume of DMSO in the presence of 1 mM NADPH. Both samples were coupled to azide-biotin via click chemistry, and proteins were purified on streptavidin beads, digested with trypsin (on-bead-digest, OBD), and analyzed by MS. The total MS signal intensity (y-axis) was plotted against the distribution (x-axis), which was calculated as the ratio between spectral intensities of the DB069 sample divided by the sum of DB096 and the click control (CC) sample (DB069/(DB069+CC)) for n=1 replicate. Highlighted are all P450 enzymes (red) and other proteins (grey). **(c)** Shotgun proteomics of the input sample. Mouse liver microsomes were digested with trypsin (in-solution-digest, ISD) and analyzed by MS. Proteins were ranked on their total MS spectral intensity and the P450s are highlighted (red) with those detected and enriched in (b) show as solid lines. *, detected by on-bead digest in (b).

To identify the labeled P450s from mouse liver microsomes, we incubated mouse liver microsomes with and without 10 μM DB096 in the presence of 1 mM NADPH and performed click chemistry with azide-biotin. Biotinylated proteins were purified, digested with trypsin/Lys-C (on-bead-digest, OBD) and analysed by LC-MS/MS. Remarkably, in this one replicate, we detected 40 mouse P450s that are >2 fold more abundant upon DB096 labeling when compared to the click control (**Figure 5b and Supplemental File S4**).

To investigate the abundance of P450s in our input samples, we also performed in-solution-digest of the liver microsomes with trypsin/Lys-C and analyzed the global protein content by LC-MS/MS. Amongst the 1410 detected mouse proteins, we detected 48 mouse P450s (**Figure 5c** and Supplemental **File S5**). A remarkable 20 P450s are in the top 150 most abundant proteins, with Cyp2c29 at rank-1 (**Figure 5b-c**). The high P450 content in mouse liver microsomes is consistent with being a highly active organ in detoxification of metabolites and explains why labeling of endogenous P450s is so effective in mouse liver microsomes.

### Mouse Cyp1a2-GFP active upon expression by agroinfiltration

To investigate if also a mouse P450 can be active in our transient expression system, we cloned codon-optimized mouse (*Mus musculus*) Cyp1a2 into a binary expression vector for *in-planta* expression by agroinfiltration. *Mm*Cyp1a2 was chosen because it was robustly detected upon labeling by DB096 (**Figure 5b**) and has a moderate abundance in mouse liver microsomes (rank-39, **Figure 5c**), suggesting an efficient labeling of this enzyme by DB096.

We expressed *Mm*Cyp1a2 as a fusion protein with a C-terminal GFP by agroinfiltation. Confocal imaging demonstrates that *Mm*Cyp1a2 colocalizes with RFP-HDEL in the ER network (**Figure 6a**), with a high Pearson’s correlation (0.881±0.073, Supplemental **Figure S1b**). To test if *Mm*Cyp1a2 is also active, we isolated microsomes from *N. benthamiana* leaves transiently expressing *Mm*Cyp1a2-GFP and tested these for labeling with DB086 and DB096. Labeling of these microsomes displayed a clear, NADPH-dependent fluorescent signal at 82 kDa, consistent with the MW of the *Mm*Cyp1a2-GFP fusion protein (**Figure 6b**). Thus, despite being a mouse enzyme, we could express *Mm*Cyp1a2 *in planta* and show its NADPH-dependent labeling with P450 probes, demonstrating that this heterologous expression system can be used to study active P450s from other organisms, presumably relying on plant CPRs providing reductants.

**Figure 6.**
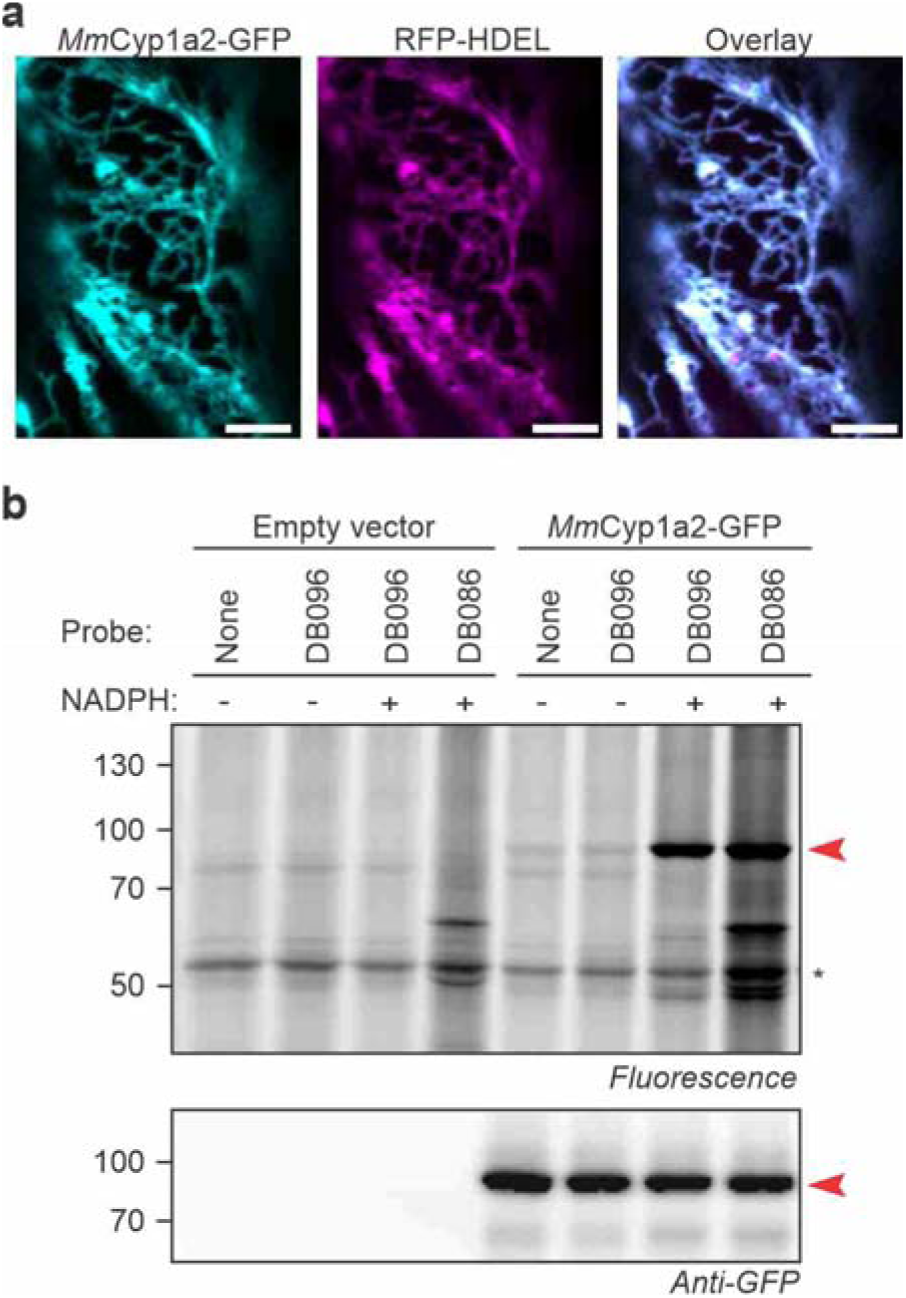
Activity-based labeling of mouse Cyp1a2 when expressed *in planta*. **(a)** *Mm*Cyp1a2-GFP localizes to the ER when expressed *in planta*. *Mm*Cyp1a2-GFP was transiently co-expressed with ER marker RFP-HDEL by agroinfiltration. Confocal images were taken 5 days after agroinfiltration. Scalebar 5 μm. Experiment was repeated twice with similar results. **(b)** NADPH-dependent labeling of mouse *Mm*Cyp1a2 expressed *in planta*. Microsomes were isolated from agroinfiltrated leaves expressing *Mm*Cyp1a2-GFP and labeled for 1.5 hr with or without 10 μM probe and with or without 1 mM NADPH. Labeled proteins were coupled to Cy2-picolyl-azide via click chemistry and visualized by in-gel fluorescence scanning. Western blot with anti-GFP antibodies shows the *Mm*Cyp1a2-GFP fusion accumulating at 82 kDa (arrowhead). *, RBSL.

### Fungal plant pathogen expresses distinct P450 activities

We next studied active P450s in *Zymoseptoria tritici*, a fungal plant pathogen that causes *Septoria tritici* blotch (STB), the most economically damaging disease of wheat in Europe [25]. Nearly 70% of the fungicides in cereal grain production in Europe are used to suppress STB, but *Z. tritici* is building resistance to these fungicides and the increased detoxification through P450s is thought to be one of the underlying mechanisms [26,27]. To identify active P450s from *Z. tritici*, we labeled microsomes isolated from *Z. tritici* cell cultures with DB096, which we used for both plant and mouse proteomics, and purified labeled proteins for LC-MS/MS analysis. A total of 13 different P450s are amongst the enriched proteins, with *Zt*CYP5078B1 being the most significantly enriched protein (**Figure 7a**, Supplemental **File S6**).

**Figure 7.**
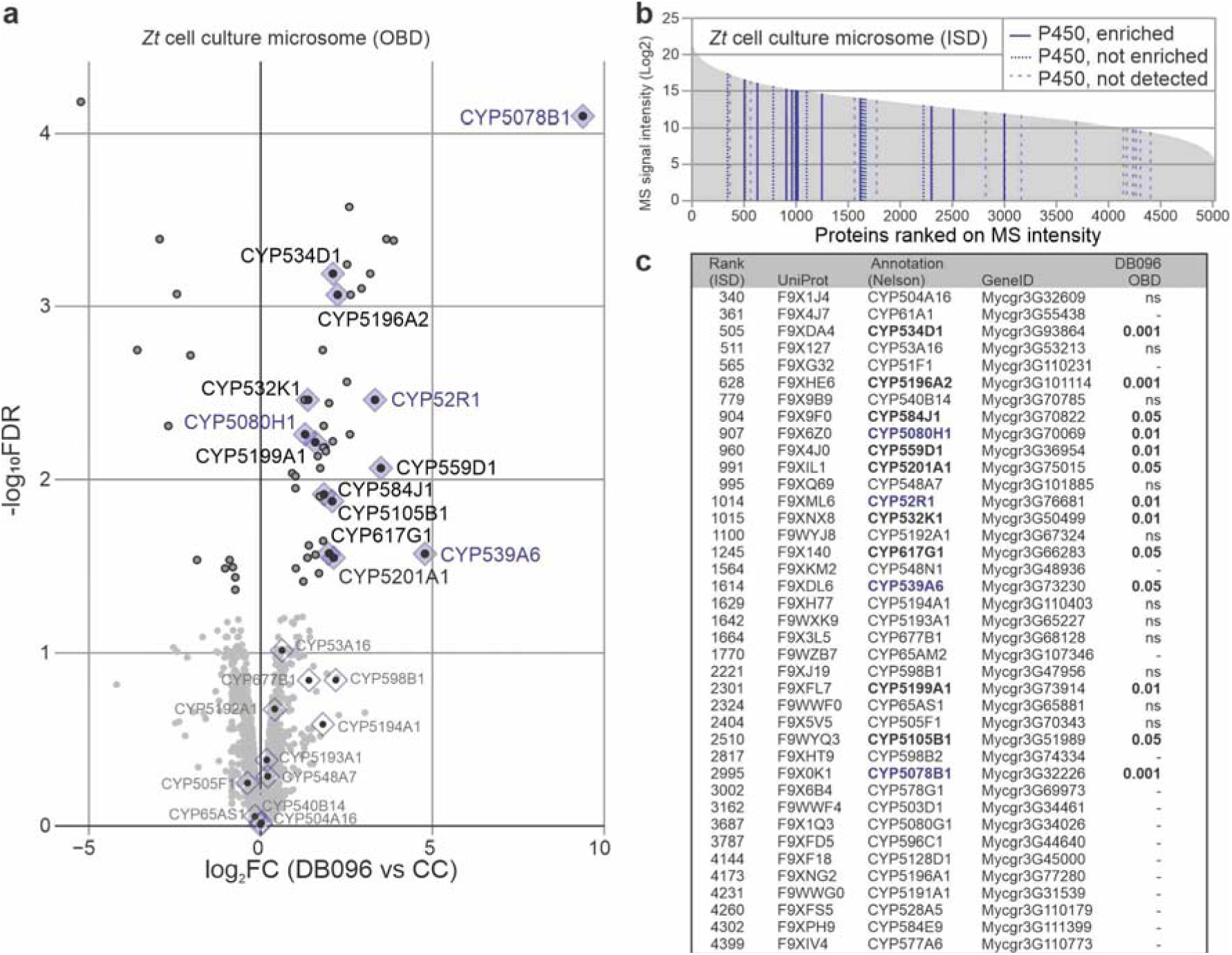
Discovery of active P450s from the fungal plant pathogen *Zymoseptoria tritici*. **(a)** Enrichment of *Zymoseptoria tritici* P450s upon labeling with DB096. Microsomes isolated from *Z. tritici* cell cultures were incubated with 30 μM DB096 or an equal volume of DMSO in the presence of 1 mM NADPH for 1h. Labelled proteins were subjected to click chemistry to attach biotin, enriched via streptavidin pull-down, and on-bead-digests (OBD) were analysed by LC-MS/MS. Volcano plots highlight proteins depleted and enriched by DB096 labelling of *Z. tritici* microsomes labelling. Enrichment is represented by the log_2_FC (P450 probe/CC) against the significance (-log_10_FDR) between the click control (CC) and DB096 using replicates. The dots indicate whether proteins were significantly enriched (black lines) or depleted (grey lines) in the DB096 sample. **(b)** Many P450s are detected in in-solution-digests (ISD) *Z. tritici* microsomes. *Z. tritici* microsomes were digested with Lys-C and trypsin and digested peptides were analysed by LC-MS/MS. All identified proteins were ranked by their spectral intensity and plotted against the spectral intensity. Lines indicate P450s that were enriched upon DB096 labeling (solid lines) or not enriched (dashed lines) or not detected (striped lines). **(c)** List of P450s detected by ISD in (b). Shown are the rank position in MS intensity, the UniProt accession code, the P450 name, the P450 name given by Donzelli, the genomic accession code and the detection of the protein upon on-bead digest (a), given with P-value if enriched, non-significant (ns) or not detected (-).

To evaluate the relative abundance of P450s in *Z. tritici* microsomes, we also performed LC-MS/MS analysis of an in solution-digest of the input samples. This analysis uncovered 40 P450s amongst the 5156 detected proteins (**Figure 7b**, Supplemental **File S7**). When ranked on intensity, the P450s are scattered across the proteome with CYP504A16 being the most abundant P450, at rank-340 (**Figures 7b and 7c**). Thus, this *Z. tritici* proteome contains many P450s at various abundance levels. Interestingly, the P450s that were enriched upon DB096 labeling are scattered across the abundance scale, indicating that not all P450s have been labeled with DB096.

### Zymoseptoria P450s have different probe reactivities

To test if also fungal P450s can be studied upon heterologous expression *in planta*, we selected P450s from four different P450 families that are amongst the DB096-enriched proteins: *Zt*CYP52R1, *Zt*CYP539A6, *Zt*CYP5078B1 and *Zt*CYP5080H1. Codon-optimized open reading frames encoding these P450s were cloned into the binary expression system fused to a C-terminal GFP. All four P450-GFP fusion proteins were detected with an anti-GFP western blot in microsomes isolated from agroinfiltrated leaves expressing these P450s (**Figure 8**). Labeling of these microsomes with the various P450 probes demonstrated that all these P450-GFP fusions can be labeled in an NADPH-dependent manner (**Figure 8**). Interestingly, however, labeling efficiencies for the probes differ per P450, demonstrating that these P450s have different labeling reactivities towards the different P450 probes.

**Figure 8.**
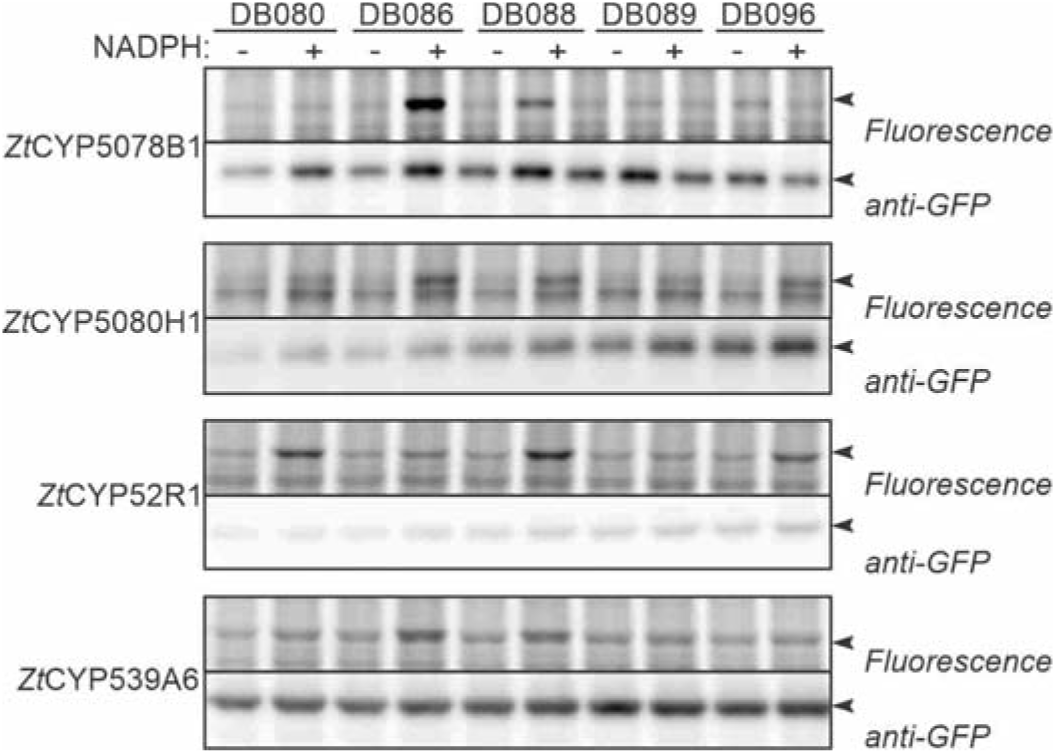
Z*y*moseptoria P450s have different reactivities to the P450 probes. Microsomes from agroinfiltrated *N. benthamiana* leaves expressing GFP-tagged *Zt*P450s were isolated and labelled with 5 μM probes in the absence or presence of 1 mM NADPH for 1h. Labelled proteins were coupled to a fluorophore via click chemistry and visualised by in-gel fluorescence scanning. Anti-GFP western blot is shown as protein expression and loading control. Similar results were obtained in two additional independent replicates.

## Discussion

We used activity-based proteomics to detect active P450s from microsomes of agroinfiltrated leaves, mouse liver and fungal cell cultures to display numerous active P450s, demonstrating a broad reach of the used chemical probes. Using transient expression of GFP-tagged P450s, we could confirm NADPH-dependent labeling of various P450s representing six P450 families specific to plants, animals and fungi, indicating that this transient expression platform can be used to characterize specific P450s further.

### Activity-based proteomics identifies a broad range of P450s in plants, mammals and fungi

We demonstrated that P450 probes developed for mammalian P450s also label fungal and plant P450s, despite representing different families. We showed the enrichment of six plant P450s representing four plant-specific P450 families [28], and 13 fungal P450s, each representing a different fungal P450 family [29]. And in addition to the five mouse P450 detected earlier in mouse liver microsomes (Cyp1a2, Cyp3a11, Cyp2c29, Cyp2d9 and Cyp2d10, [11]), our data indicate the labeling of an additional 35 P450s in mouse liver microsomes, representing an additional seven different P450 families. These data indicate that many more active P450s can be discovered with these P450 probes.

However, these probes do not label all P450s. For instance, not all P450s detected in microsomal fractions of *Z. tritici* are enriched upon DB096 labeling, indicating that they are not labeled by this probe. This includes *Zt*CYP51F1, which is an important P450 involved in azole fungicide resistance [30]. Likewise, transiently expressed fungal P450s have clearly distinct reactivities towards the different probes, indicating that some probes do not label P450s that can react with other probes. This indicates that one way to broaden the reach of P450 labeling in chemical proteomic experiments is to use probe cocktails. Some P450s, however, have very specific substrates and are unlikely to be labeled with these broad-range probes, requiring the development of specific probes.

Despite probe selectivity, our experiments indicate that the majority of the plant and fungal P450s that are available in microsomes can be labeled with the tested probes. An important limitation for detection by mass spectrometry seems to reside in the relative P450 abundance in microsomal fractions. This pattern is most obvious from that fact that of the 13 *N. benthamiana* P450s detected in microsomal fractions, only the five most abundant P450s were enriched upon labeling. This phenomenon might also explain why we detect 40 enriched P450s from mouse liver microsomes, where P450s are very abundant, and only five in leaf microsomes, where P450s are much less abundant. These observations indicate that more P450s can be detected upon enrichment from tissues with a higher abundance of P450s.

The detected active P450s may have important functions. In addition to confirming *Zm*CYP81A9 labeling, our proteomic analysis has revealed which P450s are active in agroinfiltrated *N. benthamiana*, which has become an important platform for metabolic pathway reconstruction where side reactions caused by endogenous P450s can dramatically limit yields [31]. Likewise, our proteomic analysis has also revealed which P450s are active in *Zymoseptoria tritici*, an important fungal wheat pathogen that uses P450s to detoxify fungicides.

Nevertheless, activity-based proteomics of P450s remains challenging. This method requires the isolation of microsomes to reach high P450 levels. This method is also hampered by substantial background labeling. Since background labeling also occurs in the click chemistry control, a large amount of background labeling can be avoided by developing biotinylated P450 probes that do not require click chemistry for detection. The range of detected P450s can quickly expand when using microsomes that are rich in P450s, as illustrated with mouse liver microsomes. Activity-based proteomics of P450s will be instrumental for the identification of active P450s in tissues that can e.g. detoxify agrochemicals or have established unique metabolic pathways.

### Agroinfiltration of GFP-tagged P450s enables profiling of diverse P450s

The combination of transient expression of GFP-tagged P450s with activity-based labeling is a useful innovation to facilitate further characterization of P450s. The fusion with GFP is an important detail because this facilitates both the rapid detection of protein expression before isolation of microsomes, and it shifts the molecular weight above that of endogenous P450s and RBCL, which causes strong background labeling. The ability to display active P450s on gel with activity-based probes facilitates a further characterization of P450s, e.g. to study their regulation and sensitivity to putative inhibitors, even for P450s for which substrates remain to be identified. Maize *Zm*CYP81A9 and *Z. tritici* P450s, for instance, are of interest because of important roles in detoxification of herbicides and fungicides, respectively [18,30]. The original *Zm*CYP81A9 activity assays involves reaction monitoring with mass spectrometry which requires access to analytic equipment. Imaging of fluorescence from proteins gels, however, can be done on equipment that most labs already have available. P450s from other plants are frequently expressed by agroinfiltration to reconstitute metabolic pathways in agroinfiltrated leaves [31,32]. We showed that, despite being dependent on ER-resident reductases, P450s even from animals and fungi are also active in *N. benthamiana*. Transiently expressed mouse *Mm*Cyp1a2 is correctly targeted to the ER in agroinfiltrated *N. benthamiana* and is reactive towards P450 probes, indicating that it receives reductants from *N. benthamiana* CPRs in the ER membrane.

Having established P450 profiling beyond mammals, we can now use this method to characterize P450s globally, both in P450-rich proteomes and upon overexpression by agroinfiltration. The agroinfiltration platform can now be used to detect the selective suppression of labeling by P450 inhibitors and substrates and will facilitate the development of probes and inhibitors for specific P450s. Our work therefore can provide an important foundation for future studies on P450s in diverse eukaryotic organisms.

## Materials and methods

### Chemical probes and inhibitors

The synthesis of probes DB088, DB089 and DB096 was carried out with minor alterations to the established routes as described previously [11]. Starting from the parent acids, a sequence involving i) protection as the respective methyl esters; ii) Sonogashira reactions to install the TMS-protected acetylene; and iii) global deprotection, resulted in the alkynyl acids. These acids underwent smooth conversion to DB088, DB089, and DB096 using amide coupling procedures.

The synthesis of DB086 retained the general strategy for probe synthesis with an additional step required to build the coumarin core and activate the phenolic position. In order to build the coumarin core, the Knoevenagel condensation of 2,4-hydroxybenzaldehyde with diethyl malonate was employed. From this, triflation of the phenolic position enabled the established Sonogashira reaction, deprotection and amide coupling to be carried out giving DB086.

Similarly, DB080 could be produced with only minor alterations to established conditions. Demethylation of methoxsalen was carried out with BBr3, the resulting phenol underwent smooth conversion to the requisite ester upon reaction with methyl bromoacetate. Deprotection gave access to the DB080 precursor which gave DB080 after amide coupling.

A *de novo* approach to DB143 was employed on account of stability issues faced during the Krapcho decarboxylation employed previously [11]. Propiolic acid was hydrobrominated to produce the respective (E)-vinylbromide, this was subsequently reduced with LiAlH_4_ to give the respective alcohol. This was amenable to a Sonogashira reaction with TIPS-acetylene to give the TIPS-protected enyne that was able to undergo conversion to DB143 using a route analogous to that published earlier [11], requiring only an additional deprotection step to remove the TIPS-protecting group.

For further details on probe synthesis, see **Supplemental Methods.** Aliquots of the probes are available upon request. The P450 inhibitor 1-ABT (A3940-50MG) and agrochemical benoxacor (46001-50MG) were purchased from Sigma-Aldrich. NADPH was purchased from MELFORD (N20140-0.1).

#### Molecular cloning

***–*** Plasmids and Primers are summarized in Supplemental **Tables S1** and **S2,** respectively. Golden Gate Modular Cloning kit [33] and Golden Gate Modular Cloning Toolbox for plants [34] were used for cloning. The sequence encoding mouse *Mm*Cyp1a2 and maize *Zm*CYP81A9 were codon-optimised for *in planta* expression and synthesized as a GeneString (GeneArt, ThermoFisher Scientific, Supplemental **Table S1**). Using BpiI restriction sites, these ORFs were first cloned into the level 0 module GG1-01, and transferred into pJK268c [35] together with GG1-55, GG1-57, GG1-70 or GG1-78 in a BsaI ligation reaction. Generated plasmids were amplified by transformation into *E. coli* and the inserts were confirmed by sequencing.

The sequences encoding *Z. tritici* P450s (CYP5078B1, CYP5080H1, CYP52R1 and CYP539A6) were amplified from *Z. tritici* complementary DNA (cDNA) using the forward and reverse primer pairs summarised in Supplemental **Table S2**. Using BsaI restriction sites, PCR products were combined with GG1-55, GG1-57, GG1-70 or GG1-78 [34] to assemble the corresponding expression plasmids. Generated expression plasmids were transformed into *E. coli* for their amplification and sequencing.

Binary plasmids were transformed into *Agrobacterium tumefaciens* strain GV3101, carrying the pMP90 helper plasmid [36]. Agrobacterium transformants were selected on LB plates (10 g/L NaCl, 10 g/L Tryptone, 5 g/L yeast extract, 15 g/L agar) containing 50 μM gentamycin, 25 μM rifampicin and 50 μM kanamycin. Single colonies were picked and grown in liquid LB (10 g/L NaCl, 10 g/L Tryptone, 5 g/L yeast extract) containing the same antibiotic used before. Glycerol stocks of the selected transformants were prepared mixing Agrobacterium cultures with 50% sterile glycerol in a 1:1 ratio.

### Plant growth and agroinfiltation

*Nicotiana benthamiana* plants (LAB accession, collected by Cleland in 1936 [37], specimen AD957110022 at State Herbarium of South Australia in Adeleide) were grown in the greenhouse at the University of Oxford on soil at 21 °C under long day conditions (16/8 h light/dark regime) and were used for agroinfiltrations when 4-5 weeks old. Use of *N. benthamiana* is compliant with the Nagoya protocol because this species was collected in 1939. Growing *N. benthamiana* is compliant with institutional, national and international guidelines and legislation since *N. benthamiana* is neither an endangered species nor at risk of extinction according to IUCN. Agrobacterium cultures (GV3101 pMP90) were grown overnight (approximately 18 h) at 28°C with agitation in LB media containing 50 μM gentamycin, 25 μM rifampicin and 50 μM kanamycin. Bacteria were collected by centrifugation at 1000 x *g* for 10 min at room temperature (RT), washed with infiltration buffer (10 mM MES, 10 mM MgCl_2_, pH 5.6 and 100 μM acetosyringone) and diluted to OD_600_= 0.5. Agrobacterium were incubated at 28 °C for 2 h and hand-infiltrated into two fully-expanded leaves using a needle-less 10 ml syringe.

### *Zm*CYP81A9 expression in yeast

The ORF of CYP81A9 was codon-optimized for expression in *Saccharomyces cerevisiae* and synthesized and cloned into pYES3 vector by TWIST Bioscience (CA, USA) and transformed into the WAT11 strain [18]. For yeast cultivation, yeast was grown in selective agar media containing adenine {6.7 g/L yeast nitrogen base without aminoacids (Sigma Y0626), 0.64 g/L complete supplement mixture minus tryptophan (CMS-Trp), 20 g/L glucose (Sigma G8270), 20 g/L bacto agar (BD Difco, 214050), 0.04 g/L adenine sulphate (Sigma A-9126)} at 30 °C for 48 h. A single yeast colony was inoculated in 25 ml of selective liquid media {6.7 g/L yeast nitrogen base without aminoacids (Sigma Y0626), 0.64 g/L complete supplement mixture minus tryptophan (CMS-Trp), 20 g/L glucose (Sigma G8270), 0.04 g/L adenine sulphate (Sigma A-9126)} and grown at 30 °C, 180 rpm, for 24 h. Next, 6 ml of the yeast culture was inoculated in 300 ml of YPGE containing adenine {10 g/L yeast extract (Sigma 92144), 10 g/L peptone (Melford P20P240), 30 ml/L ethanol, 20 g/L glucose and 0.04 g/L adenine sulphate (Sigma A-9126)} and incubated with agitation (180 rpm) at 30°C for 30 h. To induce protein expression, yeast cells were washed with water in sterile conditions, resuspended in 300 ml of YPLE containing adenine {10 g/L yeast extract (Sigma 92144), 10 g/L peptone (Melford P20P240), 30 ml/L ethanol, 20 g/L D-galactose, 0.04 g/L adenine sulphate (Sigma A-9126)} and incubated with agitation (180 rpm) at 30 °C for 16 h. Yeast cells were harvest by centrifugation and used for microsome isolation.

Yeast microsomes were prepared by washing the yeast cells in TEK buffer (50 mM Tris-HCL pH 7.5, 1 mM EDTA, 100 mM KCl). Yeast cells were resuspended in ice-cold TESB + DTT buffer (50 mM Tris-HCL pH 7.5, 1 mM EDTA, 600 mM D-Sorbitol, 30 mM DTT) and incubated for 10 min at room temperature. Afterwards, yeast cells were centrifuged at 5,000 x *g*, resuspended in ice-cold TESB (50 mM Tris-HCL pH 7.5, 600 mM D-Sorbitol) and ground using 0.5 mm glass beads and a bead mill homogenizer (BioSpec BeadBeater 1107900-105). The cell extract was filtered through a nylon cloth and centrifuged at 16,000 x *g* for 10 min. The supernatant was ultra-centrifuged at 100,000 x *g* for 1h. Pellet containing the microsomal membranes was resuspended in TEG buffer (50 mM Tris-HCL pH 7.5, 1 mM EDTA, 20% glycerol) and protein concentration was adjusted to 2 mg/ml by Bradford protein assay. Microsomes were kept at -80 °C until used.

### Protein extraction and microsomal fraction isolation from plant tissues

Agroinfiltrated *N. benthamiana* leaf tissue was collected at 4 days post-infiltration (dpi). Leaf tissue was frozen in liquid nitrogen and ground to fine powder with a mortar and pestle. Total proteins were extracted in cold homogenization buffer (100 mM potassium phosphate buffer pH 7.5, 400 mM sucrose, 1% (w/v) polyvinylpolypyrrolidone (PVPP) and 4 mM β-mercaptoethanol). The extracts were left for 20 min rotating at 4 °C for complete homogenization and centrifuged at 5,000 x *g* for 20 min at 4 °C. Supernatants containing soluble proteins were filtered through a layer of miracloth filter and ultra-centrifuged at 100,000 x *g* for 1h. Pellets containing the microsomal membranes were resuspended in resuspension buffer (100 mM potassium phosphate buffer pH 7.5 containing 20% glycerol). Total protein concentration was determined by DC protein assay Kit (Bio-Rad) with bovine serum albumin standards, protein concentration was adjusted to 0.75 mg/ml for each sample and the resulting samples were used for labelling without freeze thawing.

### Activity-based protein profiling and click chemistry

All probes were prepared at 0.5–2 mM stock solutions in dimethyl sulfoxide (DMSO). Protein samples of 50 μl were incubated with or without the indicated probes in the presence or absence of 1 mM NADPH for 1 hour at the indicated probe concentration (usually 5-10 μM). Labelling reactions were stopped by adding 1:4 (v/v) cold acetone and precipitating by 3 min centrifugation at maximum speed using a benchtop centrifuge. Protein pellets were resuspended in 44.5 μl of PBS-SDS buffer {phosphate-buffered saline (PBS) pH 7.4 with 1% (w/v) sodium dodecyl sulfate (SDS)} and heated for 5 min at 90 °C. For click chemistry, 44.5 μl of labelled proteins were treated with 2 μM Cy5 Picolyl Azide (Click Chemistry Tools, 50 μM stock in DMSO) or Fluorescein Picolyl Azide (Click Chemistry Tools, 50 μM stock in DMSO) and a premixture of 100 μM tris((1-benzyl-1H-1,2,3-triazol-4-yl) methyl)amine (TBTA; 3.4 mM stock in DMSO:t-butanol 1:4), 2 mM tris(2-carboxyethyl)phosphine hydrochloride (TCEP; 100 mM stock in water) and 1 mM copper(II) sulfate (CuSO_4_; 50 mM stock in water). Samples were incubated for 1 hour at RT in the dark and quenched by acetone precipitation. Protein pellets were resuspended in 50 μl of 2x gel loading buffer (100 mM Tris-HCl pH 6.8, 200 mM dithiotreitol, 4% SDS, 20% glycerol and 0.02% bromophenol blue) and heated for 5 min at 90 °C.

### *Zm*CYP81A9 activity assay with bentazon

Microsomes of yeast expressing CYP81A9 (17µl of 2 mg/ml) were incubated with 200 μM bentazon in the presence or absence of 1 mM NADPH and in a final volume of 100 μl of 50 mM Tris-HCl pH 7.5 buffer, at 28 °C for 2 hours. For inhibition experiments, yeast microsomes were pre-incubated with 300 μM P450 probes or 1-ABT for 10 min in the presence of 1 mM NADPH before adding 200 μM bentazon. Reactions were stopped by adding 100 μl acetonitrile: HCl (99:1) and proteins were precipitated by 3 min centrifugation at maximum speed using a benchtop centrifuge. Supernatants were diluted by adding 1:5 (v/v) acetonitrile in glass vials. Samples were analysed by LC-MS using the electrospray ionization interface (ESI) in negative mode. Bentazon and hydroxybentazon were detected at retention times of 1.57 and 1.49 minutes, respectively. Data is expressed in amount of hydroxylated bentazon detected (Peak area, AU) in the incubated microsomes.

### In-gel fluorescence scanning and western blot

Labelled proteins were loaded on 10-12% SDS-PAGE gels and separated at 150V. Fluorescence scanning of acrylamide gels was performed on an Amersham Typhoon 5 scanner (GE Healthcare Life Sciences) with Cy2 and Cy5 settings using the 488nm and 635nm lasers, respectively. Scanned gels were stained with Coomassie Brilliant Blue G-250. Image J was used for the quantification of the signals.

For western blot analysis, proteins separated in SDS-PAGE gels were transferred onto a PVDF membrane using Trans-Blot Turbo Transfer System (Bio-Rad). Membranes were blocked in 5% (w/v) milk in TBS-T (50 mM Tris-HCl pH 7.6, 150 mM NaCl and 0.1% Tween-20) overnight at 4 °C, incubated in 1:5,000 anti-GFP-HRP (Abcam, ab6663) for 1.5 hours at RT and washed in TBS-T for 5 times for 5 min and in TBS for the last wash. Clarity Western ECL substrate (Bio-Rad) was used for chemiluminescent protein detection. When performing streptavidin-HRP protein blots, samples were processed the same way except that membranes were blocked in 1% (w/v) BSA in TBS-T and incubated in 1:1,000 streptavidin-HRP (Sigma, S2438).

### Filter-aided sample preparation (FASP) for mass spectrometry

Mouse liver microsomes (purchased from Thermo: GIBCO Mouse (CD-1) Microsomes) were diluted to 1 mg/ml protein concentration. YM-10 microcon centrifugal filters (Millipore) were washed with 200 μl 0.1% trifluoroacetic acid (TFA) in 50% acetonitrile (ACN) and centrifuged at 15,000 x *g* for 15 min at room temperature. Filters were pre-equilibrated with 200 μl of 8 M urea in 100 mM triethylammonium bicarbonate (TEAB) pH 8.5 and centrifuged at 15,000 x *g* for 15 min at room temperature. Afterwards, an equivalent of 200 μg total protein were mixed with 200 μl of 8 M urea in 100 mM TEAB, pH 8.5 in the pre-treated filters. Filters were centrifuged at 15,000 x *g* for 40 min at room temperature and retained proteins were washed with 200 μl of 8 M urea in 100 mM TEAB a total of 4 times. Protein reduction and alkylation were achieved by sequential incubation of the concentrates in 200 μl of 8 M urea in 100 mM TEAB containing tris(2-carboxyethyl) phosphine hydrochloride (TCEP; 10 mM final concentration, 30 min incubation at room temperature) and 2-chloroacetamide (50 mM final concentration, 30 min incubation at room temperature in the dark) followed by centrifugation. The concentrates were washed with 6 M urea in 50 mM TEAB a total of 2 times and subjected to proteolytic digestion with 2.5 μg of lysyl endopeptidase (Lys-C from FUJIFILM Wako, in 50 mM Tris-HCL pH 8, 4 hours incubation at 37 °C) and 5 μg of trypsin (Trypsin Gold from Promega, in 50 mM TEAB pH 8.5, overnight incubation at 37 °C). Digested peptides were collected by centrifugation, and filters were washed with 150 μl 0.1% TFA and 150 μl 0.1% TFA in 50% ACN. The combined filtrates were dried by vacuum centrifugation and submitted for MS analysis.

### Large-scale labelling, affinity purification

Microsomal fractions (1.5 ml of 0.75 mg/ml) were incubated with or without the indicated probes (20 μM) in the presence or absence of 1 mM NADPH for 1 h. Labelling reactions were stopped by adding 4 volumes of ice-cold methanol, 1 volume of ice-cold chloroform and 3 volumes of water and centrifugation for 40 min (4,000 x *g*, 4 °C). Protein pellets were resuspended in 1 ml of PBS-SDS buffer by bath sonication and heated at 90 °C for 10 min. Labelled proteins were biotinylated with 30 μM azide-PEG3-biotin (Sigma-Aldrich, 2 mM stock in DMSO) using click chemistry as described above. Samples were incubated for 1 h at room temperature in the dark, quenched by the addition of 10 mM EDTA and proteins were precipitated via the methanol/chloroform method. Protein pellets were resuspended in 1.5 ml of 1.2% (w/v) SDS dissolved in PBS by bath sonication and diluted by adding 7.5 ml of PBS. The resulting solution was incubated with 100 μl of pre-equilibrated Pierce High Capacity Streptavidin Agarose beads (Thermo Fisher Scientific) for 2h at room temperature. Agarose beads containing the labelled proteins were collected by centrifugation at 1,000 g for 5 min at room temperature. Agarose beads were washed successively three times with 10 ml of PBS-SDS buffer, three times with PBS and a final wash with 10 ml of water.

### On-bead trypsin and Lys-C digestion

For on-bead digestion, the captured agarose beads containing the labelled proteins were treated with 500 μl 6 M urea dissolved in PBS and 10 mM TCEP for 15 min at 65 °C. A final concentration of 20 mM 2-chloroacetamide (400 mM stock in water) was added and left incubated for 30 min at 35 °C in the dark. Reaction was diluted by the addition of 950 μl of PBS and the supernatant was removed by centrifuging the beads at 500 x *g* for 2 min. 100 μl of Lys-C digestion solution (Walko, 5 μg of Lys-C was reconstituted in 200 μl 1 M urea, 50 mM Tris-HCl buffer pH 8) was added to the beads and left incubating at 37 °C for 3 h. 100 μl of trypsin digestion solution (Promega, 5 μl of reconstituted trypsin was dissolved in 100 μl 50 mM Tris-HCl buffer pH 8) was added to the beads and left incubating at 37 °C overnight. The supernatant containing the digested peptides was dried by vacuum centrifugation and send for MS analysis.

### LC-MS/MS and peptide identification using MaxQuant

MS samples were analysed as described as follows. The acidified tryptic peptides were desalted using C18 Tips (Thermo Fisher Scientific, 10 μL bed, 87782) and peptide samples were eluted with 0.1% formic acid and analysed on an Orbitrap Elite instrument (ThermoFisher Scientific) [38] attached to an EASY-nLC 1000 liquid chromatography system (Thermo Fisher Scientific). Samples were separated on an analytical column based on a fused silica capillary with an integrated PicoFrit emitter (New Objective) with Reprosil-Pur 120 C18-AQ 1.9 μm resin. Peptides were next separated using 140 min gradient of solvent and analysed with Orbitrap analyser (using Fourier Transform MS) at a resolution of 60,000 with the internal lock-mass option switched on [39].

RAW spectra were submitted to an Andromeda search in MaxQuant (version 1.5.3.30) using the default settings [40,41]. Label-free quantification and match between-runs were activated [24]. The MS/MS spectra data were searched against the UniProt Mus musculus reference database (55,412 entries). All searches included a contaminants database (as implemented in MaxQuant, 245 entries). Enzyme specificity was set to ‘Trypsin/P’ and the MS/MS match tolerance was set to ±0.5 Da. The peptide spectrum match FDR and the protein FDR were set to 0.01 (based on target-decoy approach) and the minimum peptide length was seven amino acids. For protein quantification, unique and razor peptides were allowed. Modified peptides were allowed for quantification with a minimum score of 40. Relative quantification of proteins between different MS runs is based exclusively on the label-free quantifications as calculated by the software MaxQuant, using the “MaxLFQ” algorithm [24].

Filtering of results was done in Perseus version 1.6.13.0 [42]. Briefly, data was filtered against contaminants and only those proteins found in at least one group in all two/three replicates were considered. Further analysis and graphs were performed using the Qfeatures and ggplot R packages. Imputed values were generated using a missing not at random (MNAR) and missing at random (MAR) mixed imputation over the whole matrix, and the fold change and adjusted P values (adj. P Val) were calculated using the fdr method over the three biological replicates. The correct clustering of the biological replicates by categorical groups was evaluated using principal component analysis (PCA) plots.

### Confocal Microscopy

Fluorescence microscopy was carried out using a Zeiss LSM 880 Airyscan confocal microscope with a C-Apochromat 40x/1.2 W Korr FCS M27 water-immersive objective. GFP and RFP fluorescence were detected with sequential line scanning. GFP fluorescence was excited with 2% laser power at 488 nm and emission detected with a PMT detector at 493 to 579 nm; while RFP fluorescence was excited at 561nm and emission detected with GaAsP detector at 582 to 754 nm. For z-stacks 16-bit images were captured at a slice interval of 1.31 μm. Colocalization of GFP and RFP signals was quantified with JaCoP2 plugin in Fiji software package in single images in selected regions of interest (ROIs), without saturation of fluorescence signals.

## Data Availability

The mass spectrometry proteomics data have been deposited to the ProteomeXchange Consortium via the PRIDE partner repository [43], with the dataset identifiers PXD035319 (*Nb*) and PXD046162 (*Zt*). Reviewer account details: (D7ITs2lF) and (TEYrq3Ju), respectively.

## Acknowledgements

We like to thank Urszula Pyzio for excellent plant care and Sarah Rodgers, Patricia Bowman and Caroline O’Brian for excellent technical support, and Prof. David Nelson (University of Tennessee) for annotating the detected P450s of *N. benthamiana* and *Z. tritici*.

## Author contributions

MFF designed and performed most of the experiments, with regular advice by KM, RD, SH, JS and RH; AW provided the initial probes; DB resynthesized the probes under guidance of MM and JB; RT performed confocal microscopy; CM performed phylogenetic analysis; MBH analyzed the bentazone LC-MS data; FK, MK and JS performed LC-MS/MS analysis; RH conceived and supervised the project; MFF and RH wrote the manuscript together.

## Additional information

### Supplementary Methods

**Figure S1** Fluorescence of P450-GFP and RFP-HDEL correlate in confocal microscopy.

**Table S1** Used plasmids

**Table S2** Used oligonucleotides

**File S1** On-bead digest of agroinfiltrated *Zm*CYP81A9 with CC/DB088 (ACE_0609-01)

**File S2** On-bead digest of agroinfiltrated *Zm*CYP81A9 with CC/DB096 (ACE_0609-01).

**File S3** In-solution digest of agroinfiltrated *Zm*CYP81A9 microsome (ACE_0609-02).

**File S4** On-bead digest of mouse liver microsomes with CC/DB096 (Advanced Proteomics Facility Oxford).

**File S5** In-solution digest of mouse liver microsome (Advanced Proteomics Facility Oxford).

**File S6** On-bead digest of *Zt* microsomes with CC/DB096 (Syngenta Proteomics Facility)

**File S7** In-solution digest of of *Zt* microsomes (Syngenta Proteomics Facility)

