## Supplementary material for "Discovery of active mouse, plant and fungal cytochrome P450s in endogenous proteomes and upon expression *in planta*": Table S1

### Supplementary Table S1

Font Farre *et al.* 'Discovery of active mouse, plant and fungal cytochrome P450s in endogenous proteomes and upon expression *in planta*.'

**Table S1.** Used plasmids

| **Plasmid** | **Description** | **Reference** |
|---|---|---|
| pJK268c | pL1V2-P19-F2, Binary vector | Kourelis et al., 2020 |
| GG1-01 | pL0V-SC1-15456, Level 0 Cloning vector, EC15456. | Engler et al., 2014 |
| GG1-55 | pL0M-T-35S-1-41414, Level 0, 35S term., pICH41414 | Engler et al., 2014 |
| GG1-57 | pL0M-PU-35S-TMV-3-51288, 2x35S prom. +TMVΩ, pICH51288. | Engler et al., 2014 |
| GG1-78 | pL0M-C2-eGFP-15095, Level 0 Module, eGFP for C terminal fusion. EC15095. | Engler et al., 2014 |
| pMF115 | pL2M-P19-kan-2x35S::*Mm*Cyp1a2-GFP | This work |
| pMF261 | pL2M-P19-kan | This work |
| pMF325 | pL2M-P19-kan-2x35S::*Zm*CYP81A9-GFP | This work |
| pMF330 | pL2M-P19-kan-2x35S::*Zt*CYP5078B1 | This work |
| pMF331 | pL2M-P19-kan-2x35S::*Zt*CYP5080H1 | This work |
| pMF348 | pL2M-P19-kan-2x35S::*Zt*CYP52R1 | This work |
| pMF349 | pL2M-P19-kan-2x35S::*Zt*CYP539A6 | This work |
