## Supplementary material for "Discovery of active mouse, plant and fungal cytochrome P450s in endogenous proteomes and upon expression *in planta*": Table S2

### Supplementary Table S2

Font Farre *et al.* ‘Discovery of active mouse, plant and fungal cytochrome P450s in endogenous proteomes and upon expression *in planta*.’

**Table S2** Used oligonucleotides

| **Oligo name** | **Sequence (5’-3’)** |
|---|---|
| FW_ *Zt*CYP5078B1 | TTGGTCTCAAATGGCTTTACCAGCACTTCTC |
| RV_ *Zt*CYP5078B1 | TTGGTCTCACACCGCAAGATCGCTCTCTG |
| FW_ *Zt*CYP5080H1 | TTGGTCTCAAATGTCGCTACTGACGG |
| RV_ *Zt*CYP5080H1 | TTGGTCTCACACCTACCCTTGGGCTCAC |
| FW1_ *Zt*CYP52R1 | TTGGTCTCAAATGCATAACGTCGCTCTG |
| FW2_ *Zt*CYP52R1 | TTGGTCTCAGAAACCGGTATCGATTATTCTG |
| RV1_ *Zt*CYP52R1 | TTGGTCTCATTTCCGCAGGGATGAAGATAG |
| RV2_ *Zt*CYP52R1 | TTGGTCTCACACCCTCCGCCTCGTGCAAC |
| FW1_ *Zt*CYP539A6 | TTGGTCTCAAATGCTCGTCGCCCTTATC |
| FW2_ *Zt*CYP539A6 | TTGGTCTCAGAAACCCTCCGGCTTTACC |
| RV1_ *Zt*CYP539A6 | TTGGTCTCATTTCATTCATTGTATGTGTGAGG |
| RV2_ *Zt*CYP539A6 | TTGGTCTCACACCTGACCTGCTTGCCTTC |

#### *Zm*CYP81A9

TTGAAGACAAAATGGACAAGGCTTACATTGCTGCTCTGTCTGCTGCTGCTCTTTTCCTTCTTCACTACCTGCTTGGTAGACGTGCTGGTGGTGAAGGTAAGGCTAAAGCTA
AGGGCTCTAGAAGAAGGCTGCCTCCTTCACCTCCTGCTATTCCTTTCTTGGGTCACCTCCATCTTGTGAAGGCTCCATTCCATGGTGCTCTTGCTAGACTTGCTGCTAGGCA
TGGTCCTGTGTTCTCTATGAGACTTGGAACTCGGAGGGCTGTCGTGGTTTCTTCACCTGATTGTGCTAGGGAATGCTTCACCGAGCACGATGTGAATTTCGCTAACAGGCC
TCTGTTCCCGTCTATGAGGCTTGCTTCTTTCGATGGCGCTATGCTGAGCGTGTCAAGCTATGGTCCTTATTGGAGGAACCTTAGGCGTGTTGCTGCTGTGCAACTTCTTTCA
GCTCATAGAGTGGGTTGCATGGCTCCAGCTATTGAAGCTCAGGTGAGAGCTATGGTGAGAAGGATGGATAGAGCTGCTGCAGCTGGTGGCGGTGGTGTTGCTAGAGTTC
AACTTAAGAGAAGGCTGTTCGAGCTGAGCCTGTCTGTGCTGATGGAAACTATTGCTCACACCAAGACCAGCAGAGCTGAGGCTGATGCTGATAGCGATATGTCTACTGA
GGCCCACGAGTTCAAGCAGATTGTGGATGAGCTTGTGCCTTACATCGGCACTGCTAACAGGTGGGATTACCTTCCTGTGCTGAGATGGTTCGATGTGTTCGGTGTGAGGA
ACAAGATCCTGGATGCTGTTGGTAGAAGGGATGCTTTCCTTGGCAGGCTTATCGATGGTGAGAGAAGGCGTCTTGATGCTGGTGATGAGAGCGAGAGCAAGTCAATGAT
TGCTGTGCTTCTGACCCTGCAGAAGTCTGAGCCTGAGGTTTACACCGATACCGTGATTACTGCTCTGTGCGCTAACCTTTTCGGTGCTGGTACTGAGACTACTTCTACTAC
TACCGAGTGGGCCATGAGCCTGCTTTTGAATCATAGAGAGGCACTGAAGAAGGCCCAGGCTGAAATTGATGCTGCTGTTGGAACATCTAGGCTGGTGACTGCTGATGAT
GTGCCTCACCTTACTTACCTGCAGTGCATCGTGGATGAGACTCTTAGGCTTCATCCTGCTGCTCCTCTGCTTTTGCCTCATGAATCTGCAGCTGATTGCACCGTTGGTGGTT
ACGATGTTCCTAGGGGTACTATGCTGCTTGTGAATGTGCATGCTGTGCATAGGGATCCTGCTGTGTGGGAAGATCCTGATAGATTCGTTCCCGAGAGGTTCGAAGGTGCT
GGCGGTAAGGCTGAAGGTAGACTTCTTATGCCTTTCGGCATGGGCAGACGTAAGTGTCCTGGTGAAACTCTTGCTCTTAGGACCGTGGGTCTTGTGCTTGCTACTTTGCTT
CAGTGCTTCGATTGGGATACCGTGGATGGTGCTCAAGTGGACATGAAGGCTTCAGGTGGTCTTACTATGCCTAGGGCTGTTCCTCTTGAGGCTATGTGCAGACCTAGAAC
TGCTATGAGGGGTGTGCTTAAGAGGCTGGGTGTTGTCTTCAA

#### *Mm*Cyp1a2

TTGAAGACAAAATGGCGTTCTCCCAGTACATCTCCTTAGCCCCAGAGCTGCTACTGGCCACTGCCATCTTCTGTTTAGTGTTCTGGATGGTCAGAGCCTCAAGGACCCAGG
TTCCCAAAGGCCTGAAGAATCCACCCGGACCCTGGGGCTTGCCCTTCATTGGGCACATGCTGACTGTGGGGAAGAACCCACACCTGTCACTGACACGGCTGAGTCAGCA
GTATGGGGACGTGCTGCAGATCCGCATCGGCTCCACTCCTGTGGTGGTGCTGAGCGGCCTGAACACCATCAAGCAGGCCCTGGTGAGGCAGGGAGATGACTTCAAGGGC
CGACCAGACCTCTACAGCTTCACACTTATCACTAACGGCAAGAGCATGACTTTCAACCCAGACTCTGGACCCGTGTGGGCTGCCCGCCGGCGCCTGGCCCAGGATGCCCT
GAAGAGCTTCTCCATAGCCTCGGACCCGACGTCAGCATCCTCTTGCTATTTGGAGGAGCACGTGAGCAAGGAGGCTAACCATCTCGTCAGCAAGCTTCAGAAGGCGATG
GCAGAGGTTGGCCACTTCGAACCAGTCAGCCAGGTGGTGGAATCGGTGGCTAACGTCATTGGTGCCATGTGCTTTGGGAAGAACTTCCCCCGGAAGAGCGAGGAGATGC
TGAACATCGTGAATAACAGCAAGGACTTTGTGGAGAATGTCACCTCAGGGAATGCAGTGGACTTCTTCCCGGTCCTGCGCTACCTGCCCAACCCGGCCCTCAAGAGGTTT
AAGACCTTCAATGATAACTTCGTGCTGTTTCTGCAGAAAACTGTCCAGGAGCACTACCAAGACTTCAACAAGAACAGTATCCAAGACATCACAAGTGCCCTGTTCAAGCA
CAGCGAGAACTACAAAGACAATGGCGGACTCATCCCCGAGGAGAAGATTGTCAACATTGTCAATGACATCTTTGGAGCTGGCTTTGACACAGTCACCACAGCCATCACC
TGGAGCATTTTGCTACTTGTGACATGGCCTAACGTGCAGAGGAAGATCCATGAGGAGCTGGACACGGTGGTTGGCAGGGATCGGCAACCACGGCTTTCTGACCGTCCCC
AGCTGCCATATCTAGAGGCCTTCATCCTGGAGATCTACCGATACACATCCTTTGTCCCCTTCACCATCCCCCACAGCACAACGAGGGACACCTCACTGAATGGCTTCCAC
ATTCCCAAGGAGCGCTGTATCTACATAAACCAGTGGCAGGTCAACCATGATGAGAAGCAGTGGAAAGACCCCTTTGTGTTCCGCCCAGAGCGGTTTCTTACCAATAACA
ACTCGGCCATCGACAAGACCCAGAGCGAGAAGGTGATGCTCTTCGGCTTGGGAAAGCGCCGGTGCATTGGGGAGATCCCGGCCAAGTGGGAAGTATTCCTCTTCTTAGC
CATCCTGCTGCAGCATCTGGAGTTTAGTGTGCCACCGGGTGTGAAGGTGGACCTGACACCCAACTATGGGTTGACCATGAAGCCCGGGACCTGTGAACACGTCCAGGCA
TGGCCACGCTTTTCCAAGGGTGTTGTCTTCAA
