## Supplementary material for "Discovery of active mouse, plant and fungal cytochrome P450s in endogenous proteomes and upon expression *in planta*": Figure S1

### Supplementary Figure S1

Font Farre *et al.* 'Discovery of active mouse, plant and fungal cytochrome P450s in endogenous proteomes and upon expression *in planta*.'

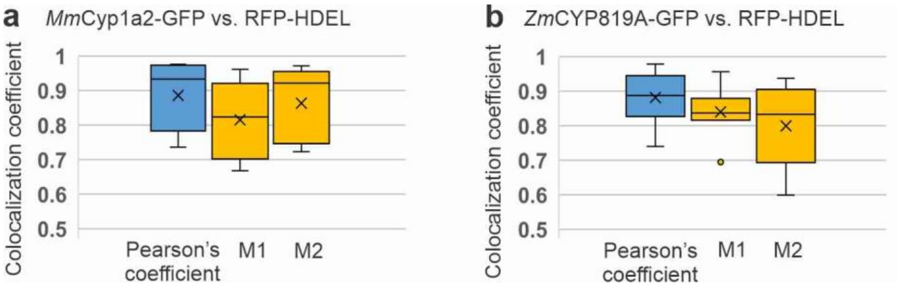


**Figure S1** Fluorescence of P450-GFP and RFP-HDEL correlate in confocal microscopy.

Colocalization of *Mm*Cyp1a2-GFP **(A)** or *Zm*CYP81A9-GFP **(B)** with RFP-HDEL was measured on a region of interest (ROI) on 10 images. Pearson's correlation coefficient (blue) and split Manders' correlation coefficients (M1 and M2, yellow) were calculated after Otsu thresholding. Shown are the mean values (x) with whiskers representing the maximum and minimum values for n=10 replicates. Signal correlation is considered strong when Pearson's coefficient value is high (above 0.8 to 1). Co-occurrence of RFP and GFP signal is tested with split Manders' coefficient M1 and M2, where M1 shows the fraction of pixels with RFP signals that overlaps with pixels with GFP signals, while M2 shows the fraction of pixels with GFP signal that overlaps with pixels with RFP signals above the threshold. M1 and M2 values range from 0 to 1, where 1 reflects perfect co-occurrence.
