## Supplemental Methods for "Discovery of active mouse, plant and fungal cytochrome P450s in endogenous proteomes and upon expression *in planta*"

### 1. Synthesis of methyl 1-bromo-2-naphthoate

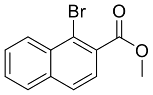


According to the modified procedure of Krätzschmar *et al*.:[2] sulfuric acid (5 drops) was added to a stirred suspension of 1-bromo-2-naphthoic acid (1.0 g, 4.0 mmol, 1.0 equiv.) and MeOH (10 mL) and the reaction mixture was heated under reflux for 18 h. The reaction mixture was cooled to rt and then solvent was removed *in vacuo* before being redissolved in EtOAc (15 mL). The organic layer was washed with sat. $NaHCO_3$ solution (20 mL) and $H_2O$ (2 × 20 mL), dried ($MgSO_4$), filtered, and the solvent removed *in vacuo* to give the title compound as a colourless oil (1.1 g, 3.9 mmol, 97%) which was used without further purification; $R_f$ 0.40 (20% EtOAc/ petroleum ether 40 – 60); $^1$H NMR (400 MHz, $CDCl_3$) δ 8.46 (ddt, *J* = 8.5, 1.5, 1.0 Hz, 1H, ArC**H**), 7.88 – 7.81 (m, 2H, 2 × ArC**H**), 7.71 – 7.57 (m, 3H, 3 × ArC**H**), 4.01 (s, 3H, C**H**$_3$); $^{13}$C NMR (101 MHz, $CDCl_3$) δ 167.9 (**C**=O), 135.2 (**C**$CO_2$Me), 132.3 (Ar**C**), 131.3 (Ar**C**), 128.6 (Ar**C**H), 128.2 (Ar**C**H), 128.1 (Ar**C**H), 127.8 (Ar**C**H), 125.8 (Ar**C**H), 122.6 (Ar**C**H), 52.7 (**C**$H_3$). One aromatic carbon resonance is unresolved. Data are in accordance with the literature.[2]

### 2. Synthesis of methyl 1-((trimethylsilyl)ethynyl)-2-naphthoate

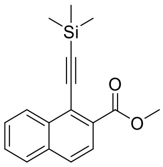


According to the modified procedure of Wright *et al*.:[1] bis(triphenylphosphine)palladium(II) chloride (0.04 g, 0.096 mmol, 2.0 mol%) was added to a solution of methyl 1-bromo-2-naphthoate (0.50 g, 1.9 mmol, 1.0 equiv.), ethynyltrimethylsilane (0.55 mL, 3.8 mmol, 2.0 equiv.), triethylamine (0.51 mL, 3.8 mmol, 2.0 equiv.), and copper(I) iodide (0.020 g, 0.23 mmol, 4.0 mol%) in MeCN (10 mL). The reaction mixture was heated to 90 °C and stirred for 16 h. The solvent was removed *in vacuo* and the resulting crude residue purified by flash column chromatography (1% EtOAc/petroleum ether 40-60 → 2% EtOAc/petroleum ether 40-60) yielding the title compound (0.19 g, 0.034 mmol, 35 %) as a brown solid. $^1$H NMR (400 MHz, $CDCl_3$) δ 8.57 – 8.52 (m, 1H, ArC**H**), 7.92 (d, *J* 8.5 Hz, 1H, ArC**H**), 7.86 – 7.82 (m, 2H, ArC**H**), 7.63 – 7.61 (m, 2H, ArC**H**), 3.99 (s, 3H, OC**H**$_3$), 0.37 (s, 9H, Si(C**H**$_3$)$_3$); *m/z* ($ES^+$) 283.3 ($M+H^+$, 100%). Data are in accordance with the literature.[1]

### 3. Synthesis of 1-ethynyl-2-naphthoic acid

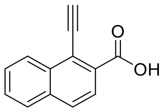

According to the modified procedure of Wright *et al*.:[1] a solution of 1 M NaOH (2 mL) was added dropwise to a stirred solution of methyl 1-((trimethylsilyl)ethynyl)-2-naphthoate (0.12 g, 0.42 mmol, 1.0 equiv.) in 2:1 EtOH:$CH_2Cl_2$ (4 mL) and then the reaction mixture was stirred at rt for 16 h. After this time, the reaction was quenched by dropwise addition of 2 M HCl (3 mL) until precipitate formed. The organics were extracted with EtOAc (3 × 8 mL), washed with brine (10 mL), dried ($MgSO_4$), filtered and the solvent removed *in vacuo* yielding the title compound as a brown solid (0.08 g, 0.41 mmol, 97%) which required no further purification. $^1$H NMR (400 MHz, acetone-$d_6$) δ 8.51 – 8.46 (m, 1H, ArC**H**), 7.98 – 7.92 (m, 2H, 2 × ArC**H**), 7.89 (d, *J* = 8.5, Hz, 1H, ArC**H**), 7.65 – 7.60 (m, 2H, 2 × ArC**H**), 4.33 (s, 1H, CC**H**); *m/z* (ESI$^-$) 195 (M-H$^+$, 100%). Data are in accordance with the literature.[1]

##### 4. Synthesis of 1-ethynyl-*N*-(hex-5-yn-1-yl)-2-naphthamide (DB089)

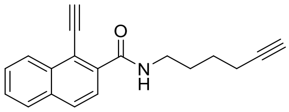

According to the modified procedure of Wright *et al*.:[1] propylphosphonic anhydride solution (≥50 wt. % in ethyl acetate, 0.12 mL, 0.36 mmol, 1.4 equiv.) was added to a stirred solution 1-ethynyl-2-naphthoic acid (0.050 g, 0.26 mmol, 1.0 equiv.), hex-5-yn-1-amine (0.040 g, 0.31 mmol, 1.2 equiv.), and DIPEA (0.18 mL, 1.0 mmol, 3.0 equiv.) in $CH_2Cl_2$ (6 mL) at room temperature. The reaction mixture was stirred at rt for 18 h. After this time, the solvent was removed *in vacuo* to give a crude residue which was further purified by column chromatography (20% EtOAc/ petrol 40-60 → 30% EtOAc/ petrol 40-60) yielding the title compound as a white solid (0.030 g, 0.052 mmol, 20%). $^1$H NMR (400 MHz, $CDCl_3$) δ 8.45 (ddt, *J* = 8.5, 1.5, 1.0 Hz, 1H, ArC**H**), 7.99 – 7.83 (m, 3H, 3 × ArC**H**), 7.68 – 7.53 (m, 2H, 2 × ArC**H**), 7.21 (br t, *J* = 5.3 Hz, 1H, CON**H**), 3.93 (s, 1H, C**H**), 3.58 (td, *J* = 7.0, 5.5 Hz, 2H, C**H**$_2$), 2.29 (td, *J* = 7.0, 2.5 Hz, 2H, C**H**$_2$), 1.98 (t, *J* = 2.5 Hz, 1H, C**H**), 1.89 – 1.75 (m, 2H, C**H**$_2$), 1.74 – 1.67 (m, 2H, C**H**$_2$); $^{13}$C NMR (101 MHz, $CDCl_3$) δ 167.1 (**C**=O), 136.4 (**C**CONH), 133.7 (Ar**C**), 133.3 (Ar**C**), 129.6 (Ar**C**H), 128.3 (Ar**C**H), 127.7 (Ar**C**H), 127.7 (Ar**C**H), 126.9 (Ar**C**H), 125.6 (Ar**C**H), 116.1 (**C**C≡CH), 89.1 (CC≡**C**H), 84.1 (C≡**C**H), 80.0 (C**C**≡CH), 68.8 (C≡**C**H), 39.7 (**C**H$_2$), 28.5 (**C**H$_2$), 25.9 (**C**H$_2$), 18.2 (**C**H$_2$); LRMS *m/z* (ES$^+$) 276.4 (M+H$^+$, 100%). Data are in accordance with the literature.[1]

##### 6. Synthesis of methyl 6-((trimethylsilyl)ethynyl)-2-naphthoate

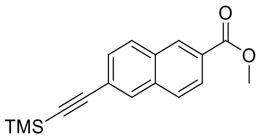

According to the modified procedure of Wright *et al.*:[1] bis(triphenylphosphine)palladium(II) chloride (0.040 g, 0.096 mmol, 5.0 mol%) was added to a solution of methyl 6-bromo-2-naphthoate (0.50 g, 1.9 mmol, 1.0 equiv.), ethynyltrimethylsilane (0.55 mL, 3.8 mmol, 2.0 equiv.), triethylamine (0.51 mL, 3.8 mmol, 2.0 equiv.), and CuI (0.020 g, 0.2 mmol, 10 mol%) in MeCN (10 mL). The reaction mixture was heated to 90 °C and stirred for 16 h. The reaction mixture was cooled to rt and the solvent was removed *in vacuo* to give a crude residue which was further purified by flash column chromatography (1% EtOAc/petroleum ether 40-60→ 2% EtOAc/petroleum ether 40-60) giving the title compound (0.51 g, 1.8 mmol, 94%) as an off white solid. $^{1}$H NMR (400 MHz, $CDCl_3$) δ 8.57 – 8.55 (m, 1H, Ar**H**), 8.06 (dd, *J* = 8.5 1.5 Hz, 1H, Ar**H**), 8.03 – 8.00 (m, 1H, Ar**H**), 7.87 (d, *J* = 8.5 Hz, 1H, Ar**H**), 7.84 – 7.80 (m, 1H, Ar**H**), 7.56 (dd, *J* = 8.5, 1.5 Hz, 1H, Ar**H**), 3.98 (s, 3H, C**H**$_3$), 0.29 (s, 9H, Si(C**H**$_3$)$_3$); $^{13}$C NMR (101 MHz, $CDCl_3$) δ 167.1 (**C**=O), 135.0 (**C**$CO_2$Me), 132.0 (Ar**C**), 131.7 (Ar**C**H), 130.8 (Ar**C**H), 129.4 (Ar**C**H), 129.3 (Ar**C**H), 128.1 (Ar**C**), 128.0 (Ar**C**H), 126.0 (Ar**C**H), 123.0 (**C**C≡C), 104.9 (**C**≡CSi), 96.3 (C≡**C**Si), 52.4 (**C**$H_3$), 0.0 (Si(**C**$H_3$)$_3$); LRMS *m/z* ($ESI^+$) 283 ($M+H^+$, 100%). Data are in accordance with the literature.[3]

### 7. Synthesis of 6-ethynyl-2-naphthoic acid

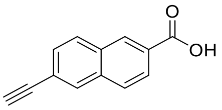


According to the procedure of Wright *et al.*:[1] a solution of 1 M NaOH (4 mL) was added dropwise to a stirred solution of methyl 6-((trimethylsilyl)ethynyl)-2-naphthoate (0.20 g, 0.71 mmol, 1.0 equiv.) in 2:1 EtOH:$CH_2Cl_2$ (8 mL). The reaction mixture was stirred at rt for 16 h before quenching by dropwise addition of HCl (6 mL) until the title compound precipitates. The aqueous layer was extracted with EtOAc (3 × 15 mL) and the organic layers combined, washed with brine (20 mL), dried ($MgSO_4$), filtered and the solvent removed *in vacuo* yielding the title compound as an off white solid (0.12 g, 0.68 mmol, 96%) which required no further purification. $^{1}$H NMR (400 MHz, acetone-$d_6$) δ 8.67 (dt, *J* = 1.5, 0.5 Hz, 1H, ArC**H**), 8.18 (d, *J* = 1.5 Hz, 1H, ArC**H**), 8.17 – 8.10 (m, 2H, 2 × ArC**H**), 8.04 (dq, *J* = 8.5, 0.5 Hz, 1H, ArC**H**), 7.65 (dd, *J* = 8.5, 1.5 Hz, 1H, ArC**H**), 3.86 (s, 1H, CC**H**); LRMS *m/z* ($ESI^-$) 195 ($M-H^+$, 100%). Data are in accordance with the literature.[3]

### 8. Synthesis of 6-ethynyl-*N*-(hex-5-yn-1-yl)-2-naphthamide (DB096)

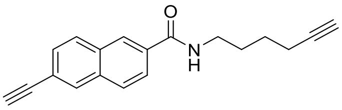


According to the modified procedure of Wright *et al.*:[1] propylphosphonic anhydride solution (≥50 wt. % in ethyl acetate, 0.23 mL, 0.77 mmol, 1.3 equiv.) was added to a stirred solution 6-ethynyl-2-naphthoic acid (0.10 g, 0.51 mmol, 1.0 equiv.), hex-5-yn-1-amine (0.080 g, 0.61 mmol, 1.1 equiv.), and DIPEA (0.25 mL, 1.5 mmol, 3.0 equiv.) in $CH_2Cl_2$ (6 mL) at room temperature. The reaction mixture was stirred for 18 h at rt. After this time the solvent was removed *in vacuo* to give a crude residue which was further purified by flash column chromatography (20% EtOAc/petroleum ether 40-60 → 30% EtOAc/petroleum ether 40-60) to give the title compound (0.060 g, 0.22 mmol, 43%) as a white solid. $^1$H NMR (500 MHz, $CDCl_3$) δ 8.23 (d, *J* = 1.5 Hz, 1H, Ar**H**), 8.02 (d, *J* = 1.5 Hz, 1H, Ar**H**), 7.87 – 7.81 (m, 3H, 3 × Ar**H**), 7.56 (dd, *J* = 8.5, 1.5 Hz, 1H, Ar**H**), 6.41 (br t, *J* = 5.5 Hz, 1H, N**H**), 3.54 (td, *J* = 7.0, 5.5 Hz, 2H, NHC$\mathbf{H_2}$), 3.20 (s, 1H, ArC≡C**H**), 2.27 (td, *J* = 7.0, 2.5 Hz, 2H, C$\mathbf{H_2}$C), 1.98 (t, *J* = 2.5 Hz, 1H, $CH_2$C≡C**H**), 1.79 (tt, *J* = 7.5, 6.5 Hz, 2H, $NHCH_2C\mathbf{H_2}$), 1.70 – 1.60 (m, 2H, $C\mathbf{H_2}CH_2C$); $^{13}$C NMR (126 MHz, $CDCl_3$) δ 167.3 (**C**=O), 134.1 (Ar**C**), 132.8 (Ar**C**), 132.2 (Ar**C**), 132.0 (Ar**C**H), 129.5 (Ar**C**H), 129 (Ar**C**H), 128.3 (Ar**C**H), 127.1 (Ar**C**H), 124.4 (Ar**C**H), 121.3 (Ar**C**), 84.1 (Ar**C**≡CH), 83.6 ($CH_2$**C**≡CH), 78.6 (ArC≡**C**H), 68.9 ($CH_2$C≡**C**H), 39.7 (NH**C**$H_2$), 28.7 $NHCH_2\mathbf{C}H_2$, 25.8 ($\mathbf{C}H_2CH_2C$), 18.2 (**C**$H_2$C); LRMS *m/z* ($ESI^+$) 276.4 ($M+H^+$, 100%); HRMS *m/z* ($ESI^+$): found 276.1383, $C_{19}H_{18}ON$ ($M+H^+$) requires 276.1383. Data are in accordance with the literature.[1]

### 9. Synthesis of methyl 4-bromobenzoate

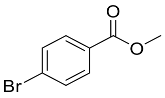


According to the procedure of Galán *et al.*:[4] Sulfuric acid (5 drops) was added to a stirred solution of 4-bromobenzoic acid (1.0 g, 5.0 mmol, 1.0 equiv.) in MeOH (13 mL) and the reaction mixture was heated to reflux for 18 h. After this time, the reaction mixture was cooled to rt and the solvent was removed *in vacuo*. The resulting crude residue was resuspended in EtOAc (15 mL) and washed with sat. $NaHCO_3$ (20 mL), $H_2O$ (2 × 20 mL), dried ($MgSO_4$), filtered and then the solvent removed *in vacuo* to give the title compound (1.1 g, 5.0 mmol, 99%) as a white solid which required no further purification. $^1$H NMR (400 MHz, $CDCl_3$) δ 7.86 – 7.79 (m, 2H, 2 × ArC**H**), 7.54 – 7.47 (m, 2H, 2 × ArC**H**), 3.84 (s, 3H, $CO_2C\mathbf{H_3}$); $^{13}$C NMR (101 MHz, $CDCl_3$) δ 166.4 (**C**=O), 131.7 (Ar**C**H), 131.1 (Ar**C**H), 129.1 (Ar**C**), 128.0 (Ar**C**), 52.3 ($CO_2\mathbf{C}H_3$). Data are in accordance with the literature.[5]

### 10. Synthesis of methyl 4-((trimethylsilyl)ethynyl)benzoate

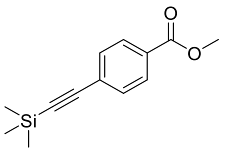


According to the modified procedure of Wright *et al.*: bis(triphenylphosphine)palladium(II) chloride (0.040 g, 0.096 mmol, 2.0 mol%) was added to a solution of methyl 4-bromobenzoate (0.41 g, 1.9 mmol, 1.0 equiv.), ethynyltrimethylsilane (0.55 mL, 3.8 mmol, 2.0 equiv.), triethylamine (0.51 mL, 3.8 mmol, 2.0 equiv.), and copper(I) iodide (0.020 g, 0.23 mmol, 4.0 mol%) in MeCN (10 mL). The reaction mixture was heated to 90 °C and stirred for 16 h. The solvent was removed *in vacuo* and the resulting crude residue purified by flash column chromatography (1% EtOAc/petroleum ether 40-60 → 2%

EtOAc/petroleum ether 40-60) giving the title compound (0.19 g, 0.034 mmol, 93 %) as an off-white solid. $^{1}$H NMR (400 MHz, $CDCl_3$) δ 8.01 – 7.92 (m, 2H, 2 × Ar**H**), 7.55 – 7.47 (m, 2H, 2 × Ar**H**), 3.91 (s, 3H, $CO_2C$**$H_3$**), 0.26 (s, 9H, Si(C**$H_3$**)$_3$); $^{13}$C NMR (101 MHz, $CDCl_3$) δ 166.4 (**C**$O_2CH_3$), 131.8 (Ar**C**H), 129.6 (Ar**C**), 129.3 (Ar**C**H), 127.7 (Ar**C**), 104.0 (**C**≡CSi), 97.6 (C≡**C**Si), 52.1 ($CO_2$**C**$H_3$), -0.2 Si(**C**$H_3$)$_3$; LRMS *m/z* ($ESI^+$) 233.1 ($M+H^+$, 100%) Data are in accordance with the literature.[6]

### 11. Synthesis of 4-ethynylbenzoic acid

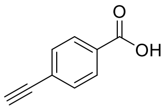

According to the modified procedure of Wright *et al.*:[1] lithium hydroxide hydrate (0.32 g, 7.5 mmol, 5.0 equiv.) was added to a stirred solution of 4-((trimethylsilyl)ethynyl)benzoic acid (0.35 g, 1.5 mmol, 1.0 equiv.) in THF (10 mL) and $H_2O$ (5 mL) and the reaction mixture was vigorously stirred at rt for 18 h. After this time, the THF was removed *in vacuo* and the remaining aqueous layer was acidified by addition of pH 1 sulfate buffer. The aqueous phase was extracted with EtOAc (3 × 10 mL), dried ($MgSO_4$), filtered and the solvent removed *in vacuo* to give the title compound (0.22 g, 1.0 mmol, 67%) as a bronze coloured solid which required no further purification. $^{1}$H NMR (400 MHz, $CDCl_3$) δ 7.85 (d, $J$ = 6.5 Hz, 2H, 2 × Ar**H**), 7.47 – 7.36 (m, 2H, 2 × Ar**H**), 3.14 (s, 1H, C≡C**H**); $^{13}$C NMR (101 MHz, $CDCl_3$) δ 167.7 (**C**=O), 131.8 (Ar**C**H), 131.0 (Ar**C**), 129.5 (Ar**C**H), 126.3 (Ar**C**), 82.8 (C≡CH), 80.0 (C≡CH); LRMS *m/z* ($ESI^-$) 145.1 ($M-H^+$, 100%). Data are in accordance with the literature.[7]

### 12. Synthesis of 4-ethynyl-*N*-(hex-5-yn-1-yl)benzamide (DB096)

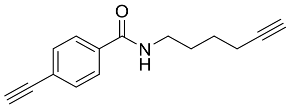

Propylphosphonic anhydride solution (50% in EtOAc, 0.25 mL, 1.0 mmol, 1.4 equiv.) was added to a stirred solution of 4-ethynylbenzoic acid (0.10 g, 0.68 mmol, 1.0 equiv.), hex-5-yn-1-amine hydrochloride (0.11 g, 0.82 mmol, 1.2 equiv.) and *N*,*N*-diisopropylethylamine (0.38 mL, 2.4 mmol, 3.0 equiv.) in $CH_2Cl_2$ (6 mL) and the reaction mixture was stirred for 1h. After this time, the solvent removed *in vacuo* and the resulting crude residue was purified by flash column chromatography (10% EtOAc/ petroleum ether 40-60 → 30% EtOAc/ petroleum ether 40-60) giving the title compound (0.096 g, 0.43 mmol, 63%) as a white solid. $^{1}$H NMR (500 MHz, $CDCl_3$) δ 7.77 – 7.71 (m, 2H, 2 × ArC**H**), 7.58 – 7.52 (m, 2H, 2 × ArC**H**), 6.31 (d, $J$ = 6.0 Hz, 1H, N**H**), 3.50 (td, $J$ = 7.0, 5.5 Hz, 2H, NHC**$H_2$**), 3.21 (s, 1H, ArC≡C**H**), 2.28 (td, $J$ = 7.0, 2.5 Hz, 2H, C**$H_2$**C), 2.00 (t, $J$ = 2.5 Hz, 1H, $CH_2$C≡C**H**), 1.82 – 1.72 (m, 2H, NHC$H_2$C**$H_2$**), 1.69 – 1.59 (m, 2H, C**$H_2$**C); $^{13}$C NMR (126 MHz, $CDCl_3$) δ 166.8 (**C**=O),

134.7 (Ar**C**), 132.3 (Ar**C**H), 126.9 (Ar**C**H), 125.3 (Ar**C**), 84.0 (CH2**C**), 82.8 (Ar**C**≡CH), 79.5 (ArC≡**C**H), 68.9 ($CH_2C$≡**C**H), 39.6 (NH**C**$H_2$), 28.6 (NH$CH_2$**C**$H_2$), 25.7 (**C**$H_2CH_2C$), 18.1 (**C**$H_2C$); LRMS *m/z* (ESI+) 248.0 (M+Na+, 100%); HRMS *m/z* (ESI+): found 226.1227, $C_{15}H_{16}ON$ ($M+H^+$) requires 226.1226. Data are in accordance with the literature.[1]

### 13. Synthesis of ethyl 7-hydroxy-2-oxo-*2H*-chromene-3-carboxylate

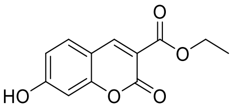


According to procedure of Vieira *et al*.:[8] piperidine (5 drops) was added to a stirred suspension of 2,4-dihydroxybenzaldehyde (0.10 g, 0.72 mmol, 1.0 equiv.) in diethyl malonate (0.23 g, 1.4 mmol, 2.0 equiv.) and the reaction mixture was stirred at rt for 18 h. The reaction mixture was acidified by addition of 2 M HCl (2 mL) until a precipitate formed. The solid was filtered and washed with $Et_2O$ (3 × 10 mL) to give a crude residue that was further purified by flash column chromatography (70% EtOAc/$CH_2Cl_2$) giving the title compound as a pale yellow solid (0.090 g, 0.74 mmol, 53 %); $^1$H NMR (400 MHz, DMSO-$d_6$) δ 11.08 (s, 1H, O**H**), 8.67 (s, 1H, C**H**CC$O_2$), 7.75 (d, *J* = 8.5 Hz, 1H, C**H**CCH), 6.84 (dd, *J* = 8.5, 2.5 Hz, 1H, CHC**H**COH), 6.72 (d, *J* = 2.0 Hz, 1H, HOCC**H**), 4.26 (q, *J* = 7.0 Hz, 2H, C**H**$_2$), 1.29 (t, *J* = 7.0 Hz, 3H, C**H**$_3$); LRMS *m/z* ($ESI^+$) 257.0 ($M+Na^+$, 100%). Data are in accordance with the literature.[8]

### 14. Synthesis of ethyl 2-oxo-7-(((trifluoromethyl)sulfonyl)oxy)-*2H*-chromene-3-carboxylate

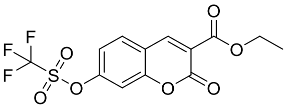


According to procedure of Starčević *et al*.:[9] triflic anhydride (0.74 mL, 4.2 mmol, 1.2 equiv.) was added dropwise to a stirred solution of ethyl 7-hydroxy-2-oxo-*2H*-chromene-3-carboxylate (0.83 g, 3.5 mmol, 1.0 equiv.) and pyridine (0.33 mL, 4.2 mmol, 1.2 equiv.) in $CH_2Cl_2$ (25 mL) cooled to 0 °C and the reaction mixture was stirred for 0.5 h. The reaction mixture was quenched by addition of $H_2O$ (30 mL). The layers were separated and the aqueous layer was extracted with $CH_2Cl_2$ (3 × 10 mL), the organic layers were combined, dried ($MgSO_4$), filtered and solvent removed *in vacuo* to give a crude residue which was further purified by flash column chromatography (20% EtOAc/ petroleum ether 40-60) to give the title compound (0.52 g, 1.4 mmol, 41%) as a yellow solid; $^1$H NMR (400 MHz, $CDCl_3$) δ 8.52 (s, 1H, C**H**CC$O_2$), 7.72 (d, *J* = 8.5 Hz, 1H, OCCHC**H**), 7.32 – 7.29 (m, 1H, OCC**H**CH), 7.28 (d, *J* = 6.0 Hz, 1H, OCC**H**), 4.43 (q, *J* = 7.0 Hz, 2H, C**H**$_2$), 1.42 (t, *J* = 7.0 Hz, 3H, $CH_2$); LRMS *m/z* ($ESI^+$) 367.3 ($M+H^+$, 100%). Data are in accordance with the literature.[9]

### 15. Synthesis of ethyl 2-oxo-7-((trimethylsilyl)ethynyl)-*2H*-chromene-3-carboxylate

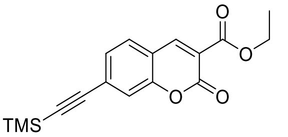


According to the modified procedure of Wright *et al.*:[1] bis(triphenylphosphine)palladium(II) chloride (0.038 g, 0.055 mmol, 5.0 mol%) was added to a solution of ethyl 2-oxo-7-(((trifluoromethyl)sulfonyl)oxy)-2*H*-chromene-3-carboxylate (0.40 g, 1.1 mmol, 1.0 equiv.), ethynyltrimethylsilane (0.31 mL, 2.2 mmol, 2.0 equiv.), triethylamine (0.29 mL, 2.2 mmol, 2.0 equiv.), and copper(I) iodide (0.023 g, 0.12 mmol, 10 mol%) in MeCN (10 mL) The reaction mixture was heated to 90 °C and stirred for 16 h. After this time, the solvent was removed *in vacuo* and the resulting crude residue purified by flash column chromatography (10% EtOAc/petroleum ether 40-60 → 20% EtOAc/petroleum ether 40-60) yielding the title compound (0.39 g, 1.0 mmol, 95%) as a yellow solid. $^1$H NMR (400 MHz, $CDCl_3$) δ 8.47 (s, 1H, C**H**CC$O_2$), 7.52 (d, *J* = 8.0 Hz, 1H, ArC**H**), 7.42 – 7.33 (m, 2H, 2 × ArC**H**), 4.41 (q, *J* = 7.0 Hz, 2H, C$\mathbf{H_2}$), 1.41 (t, *J* = 7.0 Hz, 3H, C$\mathbf{H_3}$), 0.28 (s, 9H, Si(C$\mathbf{H_3}$)$_3$); *m/z* (ESI$^+$) 315 (M+H$^+$, 100%). Data are in accordance with the literature.[1]

### 16. Synthesis of 7-ethynyl-2-oxo-*2H*-chromene-3-carboxylic acid

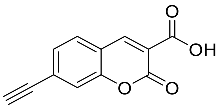


According to the modified procedure of Wright *et al.*:[1] a solution of 1 M NaOH (4 mL) was added dropwise to a stirred solution of ethyl 2-oxo-7-((trimethylsilyl)ethynyl)-2*H*-chromene-3-carboxylate (0.20 g, 0.63 mmol, 1.0 equiv.) in THF (8 mL). The reaction mixture was stirred at rt for 16 h. before quenching by dropwise addition of 2 M HCl (6 mL) to precipitate the title compound. The aqueous layer was extracted with EtOAc (3 × 15 mL) and the organic layers were combined, washed with brine (20 mL), dried ($MgSO_4$), filtered and the solvent removed *in vacuo* yielding the title compound as a yellow solid (0.10 g, 0.47 mmol, 75%) which required no further purification. $^1$H NMR (400 MHz, $CD_2Cl_2$) δ 12.01 (s, 1H, COO**H**), 8.90 (d, *J* = 1.0 Hz, 1H, C**H**CC$O_2$), 7.75 (d, *J* = 8.0 Hz, 1H, ArC**H**), 7.59 (s, 1H, ArC**H**), 7.55 (dd, *J* = 8.0, 1.0 Hz, 1H, ArC**H**), 3.51 (s, 1H, CCHCHCC≡C**H**); *m/z* (ES$^-$) 213.3 (M-H$^+$, 100%). Data are in accordance with the literature.[1]

### 17. Synthesis of 7-ethynyl-*N*-(hex-5-yn-1-yl)-2-oxo-2*H*-chromene-3-carboxamide (DB086)

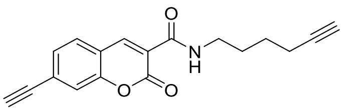

According to the modified procedure of Wright *et al*.:[1] propylphosphonic anhydride solution (≥50 wt. % in ethyl acetate, 0.10 mL, 0.35 mmol, 1.4 mmol) was added to a stirred solution of 7-ethynyl-2-oxo-2*H*-chromene-3-carboxylic acid (0.050 g, 0.23 mmol, 1.0 equiv.), hex-5-yn-1-amine (0.040 g, 0.28 mmol, 1.2 equiv.), and *N*,*N*-diisopropylethylamine (0.10 mL, 0.69 mmol, 3.0 equiv.) in $CH_2Cl_2$ (3 mL) and the reaction mixture was stirred at rt for 18 h. After this time, the solvent was removed *in vacuo* and the resulting crude residue further purified by flash column chromatography (20% EtOAc/ petroleum ether 40-60 → 30% EtOAc/ petroleum ether 40-60) giving the title compound as a white solid (0.050 g, 0.16 mmol, 70%). $^1$H NMR (400 MHz, $CDCl_3$) 8.87 (d, *J* =0.5 Hz, 1H, C**H**$CCO_2$), 8.78 (br t, *J* = 5.8 Hz, 1H, CON**H**), 7.64 (d, *J* = 8.0 Hz, 1H, ArC**H**), 7.50-7.49 (m, 1H, ArC**H**), 7.46 (dd, *J* = 8.0 Hz, 1.5, ArC**H**), 3.49 (td, *J* = 7.0, 5.8 Hz, C**H**$_2$) 3.35 (s, 1H, C**H**), 2.26 (td, *J* = 7.0, 2.5 Hz, 2H, C**H**$_2$), 1.97 (t, *J* = 2.5 Hz, 1H, C**H**), 1.80 – 1.73 (m, 2H, C**H**$_2$), 1.66 – 1.60 (m, 2H, C**H**$_2$); LRMS *m/z* ($ESI^+$) 294.3 ($M+H^+$, 100%). Data are in accordance with the literature.[1]

#### 18. Synthesis of 9-hydroxy-*7H*-furo[3,2-*g*]chromen-7-one

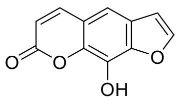

According to procedure of Shen *et al*.:[10] boron tribromide (0.19 mL, 2.0 mmol, 2.0 equiv.) was added to a vigorously stirred solution of 9-methoxy-*7H*-furo[3,2-*g*]chromen-7-one (0.22 g, 1.0 mmol, 1.0 equiv.) in $CH_2Cl_2$ (9 mL) at 0 °C, the reaction mixture was warmed to rt and stirred for 16 h. After this time, the reaction mixture was cooled to 0 °C and quenched by slow addition of saturated $Na_2CO_3$ (10 mL), the organics were extracted with EtOAc (3 × 15 mL). The organic layers were dried ($MgSO_4$), filtered, and the solvent removed *in vacuo* to give the title compound as a white solid (0.17 g, 0.85 mmol, 85%) which required no further purification. $^1$H NMR (400 MHz, DMSO-$d_6$) δ 10.65 (1H, s, O**H**), 8.12 (1H, d, *J* = 9.5, C**H**), 8.07 (1H, dd, *J* = 2.0, 1.0, ArC**H**), 7.45 (1H, d, *J* = 1.5, ArC**H**), 7.04 (1H, dd, *J* = 2.0, 0.5, ArC**H**), 6.40 (1H, d, *J* = 9.5, C**H**).

#### 19. Synthesis of methyl 2-((7-oxo-*7H*-furo[3,2-*g*]chromen-9-yl)oxy)acetate

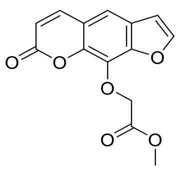

According to procedure of Wright *et al*.:[1] bromomethyl acetate (0.10 mL, 1.02 mmol, 1.2 equiv.) was added dropwise to a stirred solution of 9-hydroxy-*7H*-furo[3,2-*g*]chromen-7-one (0.17 g, 0.85 mmol, 1.0 equiv.) and potassium carbonate (0.35 g, 2.55 mmol, 3.0 equiv.) in DMF (5 mL), the reaction mixture was stirred at rt for 16 h. The solvent was removed under a stream of $N_2$ and the resulting crude residue was redissolved in EtOAc (10 mL) and washed with $H_2O$ (3 × 20 mL). The organic phase was dried ($MgSO_4$), filtered, and solvent removed *in vacuo* yielding the title compound as a white solid

(0.20 g, 0.73 mmol, 86%) which required no further purification. $^1$H NMR (500 MHz, $CDCl_3$) δ 7.79 (d, $J$ = 9.5 Hz, 1H, O=CC**H**), 7.71 (d, $J$ = 2.0 Hz, 1H, ArC**H**), 7.41 (s, 1H, ArC**H**), 6.84 (d, $J$ = 2.0 Hz, 1H, ArC**H**), 6.40 (d, $J$ = 9.5 Hz, 1H, C**H**), 5.17 (s, 2H, C**H₂**), 3.82 (s, 3H, C**H₃**); $^{13}$C NMR (126 MHz, $CDCl_3$) δ 169.3 (**C**=O), 160.1 (**C**=O), 147.2 (Ar**C**), 146.8 (Ar**C**H), 144.3 (**C**H), 142.7 (Ar**C**), 130.8 (Ar**C**), 126.1 (Ar**C**), 116.6 (Ar**C**), 114.9 (**C**H), 113.6 (Ar**C**H), 106.8 (Ar**C**H), 69.0 (**C**$H_2$), 52.3 (**C**$H_3$); LRMS *m/z* ($ESI^+$) 275 ($M+H^+$, 100%); HRMS *m/z* ($ESI^+$): found 275.0550, $C_{14}H_{11}O_6$ ($M+H^+$) requires 275.0550. Data are in accordance with the literature.[1]

### 20. Synthesis of 2-((7-oxo-*7H*-furo[3,2-g]chromen-9-yl)oxy)acetic acid

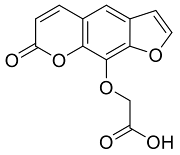

According to procedure of Wright *et al.*:[1] Lithium hydroxide (1 M in $H_2O$, 1.2 mL, 1.2 mmol, 2.0 equiv.) was added to a solution of methyl 2-((7-oxo-*7H*-furo[3,2-*g*]chromen-9-yl)oxy)acetate (0.17 g, 0.63 mmol) in THF (3 mL) and the reaction mixture stirred at rt for 16 h. After this time, the THF was removed *in vacuo* and the aqueous phase acidified with pH 1 sulfate buffer (5 mL). The organics were extracted with EtOAc (3 × 20 mL) and washed with brine (30 mL), dried ($MgSO_4$), filtered, and solvent removed *in vacuo* giving the title compound as an off white solid (0.14 g, 0.54 mmol, 85%) which required no further purification. $^1$H NMR (400 MHz, DMSO-$d_6$) 8.16 (d, $J$ = 9.5 Hz, 1H, C**H**), 8.12 (d, $J$ = 2.0 Hz, 1H, ArC**H**), 7.66 (s, 1H, ArC**H**), 7.10 (d, $J$ = 2.0 Hz, 1H, ArC**H**), 6.46 (d, $J$ = 9.5 Hz, 1H, C**H**), 5.13 (s, 2H, C**H₂**); LRMS *m/z* ($ESI^-$) 259 ($M-H^+$, 100%). Data are in accordance with the literature.[1]

### 21. Synthesis of N-(Hex-5-yn-1-yl)-2-((7-oxo-*7H*-furo[3,2-*g*]chromen-9-yl)oxy)acetamide (DB080)

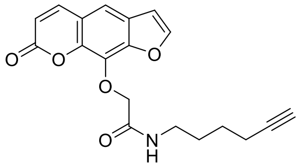

Propylphosphonic anhydride solution (≥50 wt. % in ethyl acetate, 0.15 mL, 0.52 mmol, 1.4 equiv.) was added to a stirred solution of 2-((7-oxo-*7H*-furo[3,2-*g*]chromen-9-yl)oxy)acetic acid **2.24** (0.10 g, 0.38 mmol, 1.0 equiv.), hex-5-yn-1-amine (0.050 g, 0.46 mmol, 1.2 equiv.), and DIPEA (0.50 mL, 1.0 mmol, 3.0 equiv.) in $CH_2Cl_2$ (6 mL) at room temperature. The reaction mixture was stirred overnight at rt. After this time, the reaction mixture was concentrated *in vacuo* to give a crude residue which was further purified by flash column chromatography (1% MeOH/$CH_2Cl_2$ → 2.5% MeOH/$CH_2Cl_2$) to give the title compound as a white solid (0.08 g, 0.25 mmol, 65%). $^1$H NMR (400 MHz, Acetone-$d_6$) δ 8.05 (d, $J$ = 10.0 Hz, 1H, OCOCHC**H**), 7.98 (d, $J$ = 2.0 Hz, 1H, OCHC**H**), 7.66 (s, 1H, ArC**H**), 7.64 (br, 1H, N**H**), 7.03 (d, $J$ =

2.0 Hz, 1H, OC**H**CH), 6.38 (d, *J* = 10.0 Hz, 1H, OCOC**H**CH), 4.87 (s, 2H, OC**H**$_2$), 3.37 (q, *J* = 7.0 Hz, 2H, NHC**H**$_2$), 2.31 (t, *J* = 2.5 Hz, 1H, C**H**), 2.21 (td, *J* = 7.0, 2.5 Hz, 2H, C**H**$_2$C), 1.75 – 1.63 (m, 2H, NHCH$_2$C**H**$_2$), 1.61 – 1.49 (m, 2H, C**H**$_2$CH$_2$C); $^{13}$C NMR (101 MHz, Acetone-$d_6$) δ 168.3 (**C**=O), 160.2 (**C**=O), 148.5 (OCH**C**H), 148.3 (Ar**C**O), 145.6 (OCOCH**C**H), 144.1 (Ar**C**), 131.7 (Ar**C**), 127.2 (Ar**C**), 117.7 (Ar**C**), 115.7 (Ar**C**H), 115.4 (OCO**C**HCH), 108.0 (O**C**HCH), 84.8 (**C**≡CH), 73.1 (O**C**H$_2$), 70.2 (C≡C**H**), 39.0 (NH**C**H$_2$), 29.5 (NHCH$_2$**C**H$_2$), 26.6 (**C**H$_2$CH$_2$C), 18.5 (**C**H$_2$C); LRMS *m/z* (ESI$^+$) 340.2 (M+H$^+$, 100%). Data are in accordance with the literature.[1]
